# MEX3A control of mitochondrial fitness is essential for ovarian clear cell carcinoma tumorigenesis and liver metastasis

**DOI:** 10.1101/2025.11.05.686880

**Authors:** Priyanka Vinothkumar, Pei-Yi Lin, Li-Tzu Cheng, Chen-Hsin Albert Yu, Pang-Hung Hsu, Yu-Chi Chou, Wendy W. Hwang-Verslues

## Abstract

Ovarian cancer (OC) is highly metastatic and chemoresistant. Due to heterogeneity among OC subtypes, the mechanisms underlying OC malignancy and metastasis remain largely unknown. Ovarian clear cell carcinoma (OCCC) accounts for 5-25% of OC and its incidence rate is rising. Liver metastasis is particularly high in OCCC patients and leads to significantly reduced median survival. Why OCCC metastasizes to liver at such high frequency remains elusive. We previously identified MEX3A as a key factor that promotes OCCC tumorigenesis in part by circumventing p53-mediated ferroptosis. Here, we report that MEX3A control of mitochondrial fitness, which occurs independently of p53, is essential for OCCC primary tumor growth and liver metastasis. MEX3A depletion resulted in chronic mitochondrial fragmentation and accumulation of non-functional mitochondria. MEX3A-depleted cells had decreased mitochondrial membrane potential, increased superoxide and decreased NAD^+^/NADH ratio, resulting in inhibition of oxidative phosphorylation (OXPHOS) and decreased ATP levels. In an environment enriched with mitophagy stressors, such as the liver, MEX3A-depleted OCCC cells had greatly reduced survival due to failure to recover from mitophagy. Consistent with these observations, MEX3A knockdown greatly reduced liver metastasis. Together, these data demonstrate that MEX3A-mediated mitochondrial fitness is a major factor underlying its p53-independent promotion of OCCC tumorigenesis and liver metastasis. Thus, targeting MEX3A will be a promising strategy to inhibit OCCC progression.

**Statement of significance:** Unexpected effects of MEX3A on mitochondrial fitness indicate that the need for high MEX3A expression is an OCCC vulnerability that can be exploited to uncover new treatment strategies.

## Introduction

Ovarian cancer (OC) is the deadliest gynecological cancer owing to its high metastasis, chemoresistance and recurrence (1). However, due to its complex heterogeneity, the underlying mechanisms crucial for OC malignancy remain elusive. High-grade serous ovarian carcinoma (HGSOC) is the most prevalent subtype and has been the most studied. Ovarian clear cell carcinoma (OCCC), which accounts for 5-25% of OC in different populations, is the second most prevalent subtype and its incidence rate is rising (2,3). In general, HGSOC is aggressive, has high genomic instability, and expresses mutated (mut) p53 and BRCA1/2 (4). In contrast, OCCC has wildtype (wt) p53 and low BRCA mutation frequency, but frequently is deficient in AT-rich interactive domain-containing protein 1A (ARID1A) (5). While OCCC may not be as aggressive in early stage, it generally has higher metastasis and worse prognosis in advanced stages (FIGO stages III–IV, 15% overall survival (OS) and 0% progression free survival (PFS)) compared to HGSOC (∼34% OS and 11.4% PFS) (6). Liver metastasis is particularly high in OCCC patients and these patients have significantly reduced median survival (7). The factors underlying the aggressiveness and propensity for liver metastasis of late-stage OCCC remain elusive. Numerous treatment strategies have been evaluated for OCCC, but the overall efficacy and subtype selectivity are low (8,9). Developing targeted therapies tailored to specific OC subtypes requires understanding of cancer cell intrinsic mechanisms and tumor microenvironment (TME) interactions that is still lacking for OCCC.

Bidirectional interactions between cancer cells and stromal cells are critical for tumor progression. To survive and propagate in different microenvironments during metastasis, cancer cells must tolerate various conditions, such as hypoxia, ROS, and low pH, that are different from the primary site (10). One of the key tolerance mechanisms is mitochondrial alterations. Mitochondria are the powerhouses of the cell that produce adenosine triphosphate (ATP) through oxidative phosphorylation (OXPHOS). Mitochondria are also crucial for lipid metabolism, amino acid metabolism, the tricarboxylic acid (TCA) cycle and nucleic acid metabolism (11). These essential mitochondrial metabolic functions can be altered and used by cancer cells for ATP, ROS and macromolecule production during transformation, tumor growth and metastasis. Some cancer cells can even transfer their mitochondria to stromal cells to influence their function and immune profile to impair anti-tumor immune response (12).

Mitochondrial dynamics, including fission, fusion and mitophagy, play critical roles in metabolic reprograming and mitochondrial quality control of cancer cells. Mitochondrial fission is the process where one mitochondrion divides into two or more independent organelles. It ensures proper mitochondrial partitioning and distribution during cell division, contributes to mitochondrial heterogeneity, and facilitates initiation of apoptosis (13). As a counterpart to fission, mitochondrial fusion is the process where two mitochondria physically merge and mix their contents and DNA. This process ensures a shared gene pool and optimizes overall mitochondrial function to maintain cellular health (13). A balance between fission and fusion is required for cellular homeostasis and adaptation to metabolic demands under different conditions (13,14). In cancer cells, however, the fission-fusion balance is often disrupted. Elevated mitochondrial fission typically leads to enhanced glycolysis; while increased mitochondrial fusion results in enhanced OXPHOS capacity (14). Thus, the imbalance between fission and fusion can contribute to metabolic reprogramming of cancer cells to drive tumorigenesis, metastasis and drug resistance (14). When mitochondria are damaged, mitophagy can be activated to selectively remove dysfunctional mitochondria by transporting them to lysosomes for degradation. When damaged mitochondria are removed, new and healthy mitochondria can be generated. This mitophagy-mitobiogenesis coupling is not only critical for mitochondrial quality control, but also allows cells to undergo metabolic switch between glycolysis and OXPHOS to adapt and survive under different cellular states and stresses (15). In cancer cells, mitophagy is a critical process to clear damaged or excessive mitochondria to help cancer cells adapt to dynamic and stressful microenvironments and maintain energy supply to allow tumor progression and metastasis (15). Mitochondrial dynamics also play roles in the interaction between cancer cells and microenvironment to influence tumor development and chemoresistance (16,17). For example, hypoxia can induce mitochondrial fission which contributes to cisplatin resistance in HGSOC cells (18). Also, during cancer development, malignant cells can remodel the extracellular matrix (ECM) stiffness to create a favorable microenvironment. Depending on the cell type and context, the stiff microenvironment can either promote mitochondrial fusion or induce mitochondrial fission (19,20) to facilitate cancer cell proliferation, invasion, migration, metastasis and drug resistance (21). Thus, understanding the intrinsic mitochondrial regulations in OCCC cells and the interplay between mitochondrial dynamics and the microenvironment will not only allow us to understand the biology of OCCC progression, but also provide a basis for therapeutic development to impede cancer metastasis.

Altered expression of RNA-binding proteins (RBPs) is known to promote tumorigenesis in many cancers (22), including OC (23). MEX3A is a dual functional protein which can bind RNAs via two tandemly repeated KH (KH1/KH2) RNA binding domains to regulate mRNA metabolism (24). MEX3A can also modulate protein stability via a C-terminal RING finger domain for targeted protein ubiquitination and subsequent degradation (25,26). In normal tissues, MEX3A plays a critical role in maintaining stem cell homeostasis (27,28). In cancers, MEX3A is often upregulated and its upregulation is associated with tumorigenesis and cancer progression (25,29–31). MEX3A has been found to regulate genes that enhance cancer stemness and therefore promote tumorigenesis, epithelial-mesenchymal transition (EMT) and metastasis (32,33). In OC, MEX3A upregulation has been correlated with poor prognosis (34). In HGSOC, MEX3A post-transcriptionally regulates oncogene expression to facilitate cancer cell proliferation (34). Its expression is also associated with immune infiltration based on multi-omics analyses (35).

We previously identified MEX3A as a key OCCC oncogenic factor that promotes OCCC tumorigenesis in part by circumventing p53-mediated ferroptosis (26). Here, we further report that MEX3A, independently of p53, is essential for OCCC mitochondrial fitness that contributes to tumor growth at the primary site and is crucial for liver metastasis. MEX3A depletion resulted in chronic mitochondrial fragmentation, which rendered cells unable to recover from stress-induced mitophagy. MEX3A-depleted cells had disrupted mitochondrial membrane potential, increased superoxide level, decreased NAD^+^/NADH ratio and were deficient in electron transport chain (ETC) supercomplex assembly, leading to significantly impaired OXPHOS and reduced ATP production. Consequently, MEX3A-depleted cells had impaired growth and were more sensitive to mitochondrial stress induced by different microenvironments, particularly the liver microenvironment. Consistent with this, MEX3A-depleted cells had greatly decreased liver metastasis. Together, these data demonstrate that the role of MEX3A in mitochondrial fitness is a major factor underlying its p53-independent promotion of OCCC tumorigenesis and liver metastasis. This suggests that targeting MEX3A is a promising strategy to inhibit OCCC progression.

## Materials and Methods

### Cell lines

Immortalized human endometriotic cells (IHEC; 12Z, T0764; RRID: CVCL_0Q73) was obtained from Applied Biological Materials and cultured on Type I collagen-coated plates using Prigrow III medium (TM003). Human OCCC cell line TOV21G (RRID: CVCL_3613) was purchased from the Bioresource Collection and Research Center (Taiwan), JHOC5 (RRID: CVCL_4640) and JHOC9 (RRID: CVCL_4643) from the RIKEN Bioresource Research Center (Japan), and RMG1 (RRID: CVCL_1662) from the Japanese Collection of Research Bioresources. OVCA429 (RRID: CVCL_3936) was kindly provided by Dr. Noriomi Matsumura (Kindai University, Japan). All cancer cell lines were authenticated by short tandem repeat (STR) profiling (Table S1) and tested for Mycoplasma contamination using a PCR-based assay (BIOTOOLS, TTB-GBC8). Culture conditions were as follows: TOV21G was cultured in 1:1 mixture of MCDB 105 and Medium 199 supplemented with 15% FBS, OVCA429 in DMEM with 10% FBS, JHOC9 in RPMI1640 with 10% FBS, RMG-1 in Ham’s F12 with 10% FBS and JHOC5 in DMEM/F12 with 10% FBS and 0.1mM non-essential amino acids. All media were supplemented with 1× Pen-Strep-Ampho B solution (Biological Industries, 03–033–1B). Cells were maintained at 37 °C in a humidified incubator supplemented with 5% CO₂. IHEC cells were used within five passages post-thawing, and cancer cell lines were used within 25 passages.

### Reagents

For short hairpin RNA (shRNA) experiments, shRNA lentivirus targeting MEX3A [shMEX3A#1 (TRCN230492), shMEX3A#2#2 (TRCN230494)], sh-p53#2 (TRCN342334) and the non-targeting control shRNA (shCtrl; TRC2-Scramble_ASN0003) were purchased from the National RNAi Core facility (Academia Sinica, Taiwan). Carbonyl cyanide p-trifluoromethoxyphenylhydrazone (FCCP) was purchased from MedChemExpress (HY-100410).

### Immunoblot (IB) assay

Whole-cell lysates were prepared in RIPA lysis buffer (20-188, Millipore) supplemented with 1 mmol/L PMSF (11359061001, Sigma-Aldrich), protease inhibitor cocktail (S8830, Sigma-Aldrich), and phosphatase inhibitor cocktail (TAAR-BBI3, BIOTOOLS). Lysates were sonicated on ice at 4 °C using a UP50H sonicator (Hielscher), and protein concentrations were determined using the Bradford assay (5000006, Bio-Rad). Equal amounts of protein were separated by SDS-PAGE and transferred onto PVDF membranes. Membranes were blocked with 5% skimmed milk (232100, Difco) for 1 h at room temperature (RT), followed by overnight incubation at 4 °C with primary antibodies (Table S2). After washing with PBST, membranes were incubated with HRP-conjugated anti-mouse (115-035-003; Jackson ImmunoReseaerch, RRID: AB_10015289, 1:5,000) or anti-rabbit (111-035-003, Jackson ImmunoResearch, RRID: AB_2313567, 1:5,000) secondary antibodies at room temperature for 1 h. Signals were detected using a UVP ChemStudio Plus BioImaging system, and band intensities were quantified with ImageJ software (RRID:SCR_003070).

### RNA extraction and qRT-PCR

Total RNA was extracted using TRI Reagent (T9424, Merck), and cDNA synthesis was generated using SuperScript III Reverse Transcriptase (18-080-044, Invitrogen). qRT-PCR was performed using SYBR Green 2× master mix (KM4116, KAPA Biosystems) with an Applied Biosystems QuantStudio 5 Real-Time PCR System. Relative mRNA levels were calculated using the comparative Ct (ΔΔCt) method, with cyclophilin RNA serving as the internal control. The sequences of all primers used in this study are listed in Table S3.

### Immunofluorescence (IF)

IF was performed as previously described with modifications (36). Briefly, cells on coverslips were fixed with 4% formaldehyde for 15 min at RT, permeabilized with 0.1% Triton X-100 in PBS for 15 min, and blocked with 1% BSA in PBS for 1 hour at RT. Cells were then incubated with antibodies (Table S2) overnight at 4 °C followed by goat anti-mouse or goat anti-rabbit secondary antibodies (1:200) for 1 h at 37 °C. Nuclei were stained with 4’6-diamidino-2-phenylindole (DAPI; D1306, Invitrogen, RRID: AB_2629482) and mounted using Dako mounting medium. Images were acquired using a LSM780 + ELYRA inverted confocal plus super resolution microscope, and fluorescence intensities were quantified using ImageJ software.

### RNA-sequencing (RNA-seq) analysis

Total RNA was extracted and quantified using NanoDrop (Thermo, ND 1000). All samples with OD 260/280 ratio > 1.8 and OD 230/260 ratio > 2 were sent to BIOTOOLS Company in Taiwan for sequencing. RNA quality was measured by Qsep100 Bio-Fragment Analyzer. Samples with RQN > 6.8 were used for cDNA library construction. Paired-end 150-bp reads, sequenced by NovaSeq 6000 platform, were first processed using fastp (v0.20.0, RRID:SCR_016962) to trim adapters and filter out low quality bases. The trimmed reads were then aligned to the human reference genome (GRCh38) using STAR (v=2.7.11b, RRID:SCR_004463). The Gene-level read counts were subsequently quantified using HTSeq (v2.0.4, RRID:SCR_005514). After normalization, the differential expressed genes (DEGs) between shMEX3A and shCtrl cells were determined based on an FDR *p*-value < 0.05 and an absolute log2 fold change > 1 using the R package ‘edgeR’ (v4.0.16, RRID:SCR_012802). To identify enriched functional pathways, we performed Gene Set Enrichment Analysis (GSEA) (RRID:SCR_003199) on the ranked gene list determined by their FDR *p-*value (Table S4). Expression fold change and FDR *p-*value of the enriched genes in the GSEA-OXPHOS hallmark set was listed in Table S5.

### Xenograft mouse models

All animal procedures were approved by the Institutional Animal Care and Utilization Committee of Academia Sinica, IACUC number 24-03-2145 and conducted in accordance with institutional and national guidelines. Female BALB/cAnN.Cg-Foxn1nu/CrlNarl (NUDE) (RRID: IMSR_CRL:194) mice were purchased from the National Laboratory Animal Center (Taipei, Taiwan) at 6-8 weeks of age.

#### Intra bursal injection

Pre-operative analgesia was administered via intramuscular injection of Atropine, 0.05mg/kg Atropine and subcutaneous injection of 2mg/kg Achefree. After 30 min, anesthesia was induced via intraperitoneal injection of 25 mg/kg Zoletil and 7.5 mg/kg Rompun. Once fully anesthetized, mice were placed in a lateral position with the spleen oriented upward. A small incision was made on the flank using sterile surgical tools, and the ovarian fat pad was gently exteriorized to expose the ovary. 3 × 10⁵ cells in 25 μL Matrigel (354234, Corning) were injected into the oviductal tubule leading to the ovarian bursa using an insulin syringe. The ovary was returned to the abdominal cavity, the incision was closed using sterile sutures, and antiseptic cream was applied to the wound site. Mice were monitored until recovery from anesthesia and observed daily thereafter. For TOV21G-injected mice, euthanasia was performed at 6 weeks post-injection. For JHOC5-injected mice, euthanasia was performed at 10 weeks or when tumors reached approximately 2.0 cm in diameter.

#### Intrasplenic injection

Pre-operative analgesia was administered via intramuscular injection of Atropine, 0.05mg/kg Atropine and subcutaneous injection of 2mg/kg Achefree. After 30 min, anesthesia was induced via intraperitoneal injection of 25 mg/kg Zoletil and 7.5 mg/kg Rompun. Once fully anesthetized, mice were placed in a lateral position with the spleen oriented upward. Intrasplenic injection was performed as previously described (37). A small incision was made in the left flank using sterile surgical instruments, and the spleen was gently exteriorized. 1×10⁷ cells in 50ul PBS were slowly injected directly into the splenic parenchyma using an insulin syringe without puncture through the spleen capsule to avoid leakage. Immediately after injection, a sterile, moist gauze was applied to the injection site for ∼2 min to prevent backflow of the cells. Following hemostasis, the spleen was removed using a cauterizer. The abdominal incision was closed using sterile sutures, and antiseptic cream was applied to the wound. Mice were monitored until full recovery from anesthesia and subsequently observed daily for signs of distress or complications. Mice were euthanized 4 weeks after injection.

### Liver extract preparation

Liver extract preparation was performed as previously described (38). Livers were excised from female NUDE mice and washed with ice-cold 0.05 M Tris buffer (pH 7.4) containing 0.15 M KCl. Liver tissue (∼200 mg) was minced with a sterile razor blade and homogenized using a motor and pestle in 1 mL Dulbecco’s Modified Eagle Medium (DMEM) per 200 mg tissue. The homogenate was passed through a 50 μm nylon mesh to remove fibrous tissue and centrifuged to pellet the debris. The supernatant was filtered through a 0.22 μm filter and stored at −80 °C until use. For functional assays, growth medium supplemented with 1% FBS and 10% liver homogenate was used.

### Immunohistochemistry (IHC)

Paraffin-embedded xenograft tumor sections were dewaxed in xylene, rehydrated through graded ethanol to water, and subjected to antigen retrieval using Trilogy buffer (920P-07, Cell Marque) for 10 mins under high pressure. Endogenous peroxidase activity was quenched with 3% H₂O₂ for 20 min. Sections were blocked with 5% non-fat milk in PBST for 1 h at RT and incubated overnight at 4 °C with primary antibody against CK7 (bs-1610R, Bioss, RRID: AB_10856304, 1:200) in blocking buffer. After washing, slides were incubated with HRP-conjugated rabbit polymer (K4003, Dako, Agilent Technologies; RRID: AB_2630375) for 60 min, followed by DAB visualization using the Dako REAL EnVision kit (K5007; RRID: AB_2888627) and counterstaining with hematoxylin. Images were acquired at 40× magnification using an Aperio GT450 scanner (Leica Biosystems). Positive staining intensity and area were quantified using QuPath digital analysis system (RRID: SCR_018257).

### Transmission electron microscopy (TEM)

Cells were seeded on an ACLAR embedding film (Electron Microscopy Sciences) and cultured until reaching ∼98% confluence. Monolayers were rinsed once with serum-free culture medium at RT and chemically fixed in a solution containing 2.5% glutaraldehyde, 2% paraformaldehyde, 0.1% tannic acid, and 0.1 M sodium cacodylate buffer (pH 7.2) for 30 min at RT. The fixative was prepared by mixing 2.5 ml of 10% glutaraldehyde, 1.25 ml of 16% paraformaldehyde, 0.1 ml of 10% tannic acid, 5 ml of 0.2 M sodium cacodylate buffer (pH 7.2), and 1.15 ml of distilled water to a final volume of 10 ml. Cells were washed three times for 5 min each at RT with 0.1 M sodium cacodylate buffer (pH 7.2) containing 0.2 M sucrose and 0.1% CaCl₂. Post-fixation was performed in 2% osmium tetroxide (prepared by mixing 5 ml of 4% OsO₄ with 5 ml of 0.2 M sodium cacodylate buffer, pH 7.2) for 60 min at RT. Following post-fixation, cells were washed three times in distilled water for 5 min each. En bloc staining was carried out with 1% aqueous uranyl acetate for 30 min at RT, followed by three additional washes in distilled water (5 min each). Samples were dehydrated through a graded ethanol series at RT: 30% (1 × 10 min), 50% (1 × 10 min), 70% (1 × 10 min), 90% (1 × 10 min), 95% (1 × 10 min), and 100% ethanol (3 × 10 min). Infiltration was performed sequentially with ethanol:EPON resin mixtures at the following ratios and times: 2:1 (1 h), 1:1 (1 h), 1:2 (1 h), followed by four changes of pure EPON (1 h each, with one overnight infiltration). Samples were embedded in fresh EPON resin and polymerized at 60 °C for 48 h. Ultrathin sections were prepared using an ultramicrotome and imaged with a Tecnai G2 Spirit TWIN transmission electron microscope (Thermo).

### Seahorse analysis

The Seahorse XF Mito Stress test was performed using Seahorse XF Cell Mito Stress Test Kit (Agilent 103-015-100) on a Seahorse XFe24 analyzer (Seahorse Bioscience, Agilent Technologies, Santa Clara, USA) following the manufacturer’s protocol. Total cell numbers of each well in the XF24 cell plates were used for normalization.

### Clonogenic assay

Ten thousand cells were seeded in a well of 6-well plates and cultured for 14 days. Colonies were fixed with a mixture of methanol and acetic acid (3:1) for 5 min at RT, and stained with 1% crystal violet solution (prepared in pure ethanol) for 15 min. Numbers of colonies were quantified using ImageJ software.

### Soft agar colony formation assay

Two thousand five hundred cells were mixed with a layer of 0.35% agar/complete growth medium over a layer of 0.5% agar/complete growth medium in a well of a 12-well plate. After culture at 37°C for 2-3 weeks, colonies ≥ 50 μm in diameter were counted from nine random fields per well under 40× magnification using a light microscope.

### MitoTracker staining

Cells were cultured on coverslips in complete growth media until ∼90–95% confluent. The medium was replaced with pre-warmed 1× HBSS buffer containing 100nM MitoTracker dye (M7514, Thermo Fisher Scientific; RRID: SCR_013555) and 1 μg/ml Hoechst 33342 (Hoechst, #B2261, Sigma-Aldrich) for 30 min. The images were captured by a Leica TCS SP8X WLL Confocal Super resolution microscope at 37 °C in a humidified atmosphere with 5% CO₂.

### Mitochondrial membrane potential assay

Mitochondrial membrane potential was measured using Mitochondrial Membrane Potential Assay Kit (with JC-1) (E-CK-A301, Elabscience) following the manufacturer’s protocol. Fluorescent signals were captured by an OLYMPUS IX73P2F Inverted Microscope and quantified using a VisionWorks software (Analytik Jena).

### MitoSOX assay

Superoxide levels in cells were measured using MitoSOX™ Red mitochondrial superoxide indicator (M36008, Thermo Fisher Scientific) according to the manufacturer’s instructions. Fluorescent signals were captured by an OLYMPUS IX73P2F Inverted Microscope and quantified using a VisionWorks software (Analytik Jena).

### ATP detection

ATP levels in cell lysates were measured using the ATP Detection Assay Kit – Luminescence (Cayman Chemical, Cat. No. 700410) according to the manufacturer’s instructions. Luminescence was measured using a SpectraMax® iD5 microplate reader (Molecular Devices).

### Intracellular NAD⁺/NADH measurement

The intracellular NAD⁺ and NADH were measured using the NAD⁺/NADH Colorimetric Assay Kit (WST-8) (Elabscience, Cat. No. E-BC-K804-M) following the manufacturer’s instructions and the absorbance was measured at 450 nm using a SpectraMax® iD5 microplate reader (Molecular Devices).

### Blue-Native PAGE

Mitochondrial proteins were isolated using the Mitochondria/Cytoplasmic Extraction Kit (ab65320, Abcam) following the manufacturer’s protocols, and the concentration was determined by Bradford assay. Blue-Native PAGE was performed as previously described (39). 50 μg of the extracted mitochondrial protein was mixed with a sample buffer cocktail consisting of 5 μL NativePAGE sample buffer, 4 μL 5% digitonin, and 11 μL H2O. After incubation on ice for 20 min, samples were centrifuged at 20,000 × g for 10 min at 4 °C. For each sample, 15 μL of the supernatant was collected, mixed with 1 μL Coomassie G-250 sample additive and loaded to a 3–12% gradient gel (XCell SureLock Mini-Cell system, NOVEX). After electrophoresis, proteins were transferred to PVDF membranes overnight at 4 °C using a X Cell Blot module (NOVEX). Membranes were sequentially washed in 8% acetic acid (5 min, with gentle shaking) and 100% methanol (5 min), blocked in 5% skimmed milk in PBST for 1 h, and incubated with primary antibodies (Table S2) overnight at 4 °C. Following incubation with HRP-conjugated secondary antibodies for 1 h at RT, signals were visualized using a UVP ChemStudio Plus BioImaging System, and band intensities were quantified using ImageJ software.

### Statistics

All data were presented in replicates of three or more and presented as means ± SD. Unpaired *t* test or two-way ANOVA was used to compare control and individual treatment groups. All statistical analyses were performed using Prism 10 software (RRID:SCR_002798).

### Data Availability

The RNA-seq data generated in this study are publicly available in Gene Expression Omnibus (GEO) (RRID: SCR_005012) at GSEXXX. (Due to government shutdown, submission high-throughput sequence data to GEO is currently not processed.)

## Results

### MEX3A is required for OCCC tumorigenesis and liver metastasis

A panel of OCCC cell lines including TOV21G and JHOC5, two well-established OCCC models (40,41), were used to examine MEX3A expression. Compared to immortalized human endometriotic cell line (IHEC, which served as a normal control), OCCC cell lines had higher MEX3A expression at both transcriptional and translational level (Fig. S1A and S1B). TOV21G had the highest and JHOC5 had moderately higher MEX3A expression compared to other OCCC cell lines. Because of their high MEX3A level, TOV21G and JHOC5 were used for loss-of-function assays where MEX3A was depleted with shRNA (Fig. S1C and S1D) followed by orthotopic xenograft assays. TOV21G and JHOC5 cells, with or without MEX3A depletion, were injected into the ovarian bursa of NUDE mice and inoculated for six and ten weeks, respectively. Mice were then euthanized and tumor size as well as numbers of liver nodules were evaluated. Consistent with our previous report that MEX3A is essential for OCCC tumorigenesis (26), shMEX3A cell-derived tumors were significantly smaller than those derived from shCtrl cells (Fig. 1A, 1B). MEX3A depletion also greatly decreased liver colonization compare to shCtrl cells (Fig. 1C and 1D). It is worth noting that clinical evidence has suggested that OC metastasis to distant organs, including liver, typically needs to go through hematogenous route (42). As the microenvironments of the ovary, bloodstream and liver differ from each other significantly, these observations also indicated that MEX3A expression is required for cells to tolerate and proliferate in different microenvironments.

**Figure 1.**
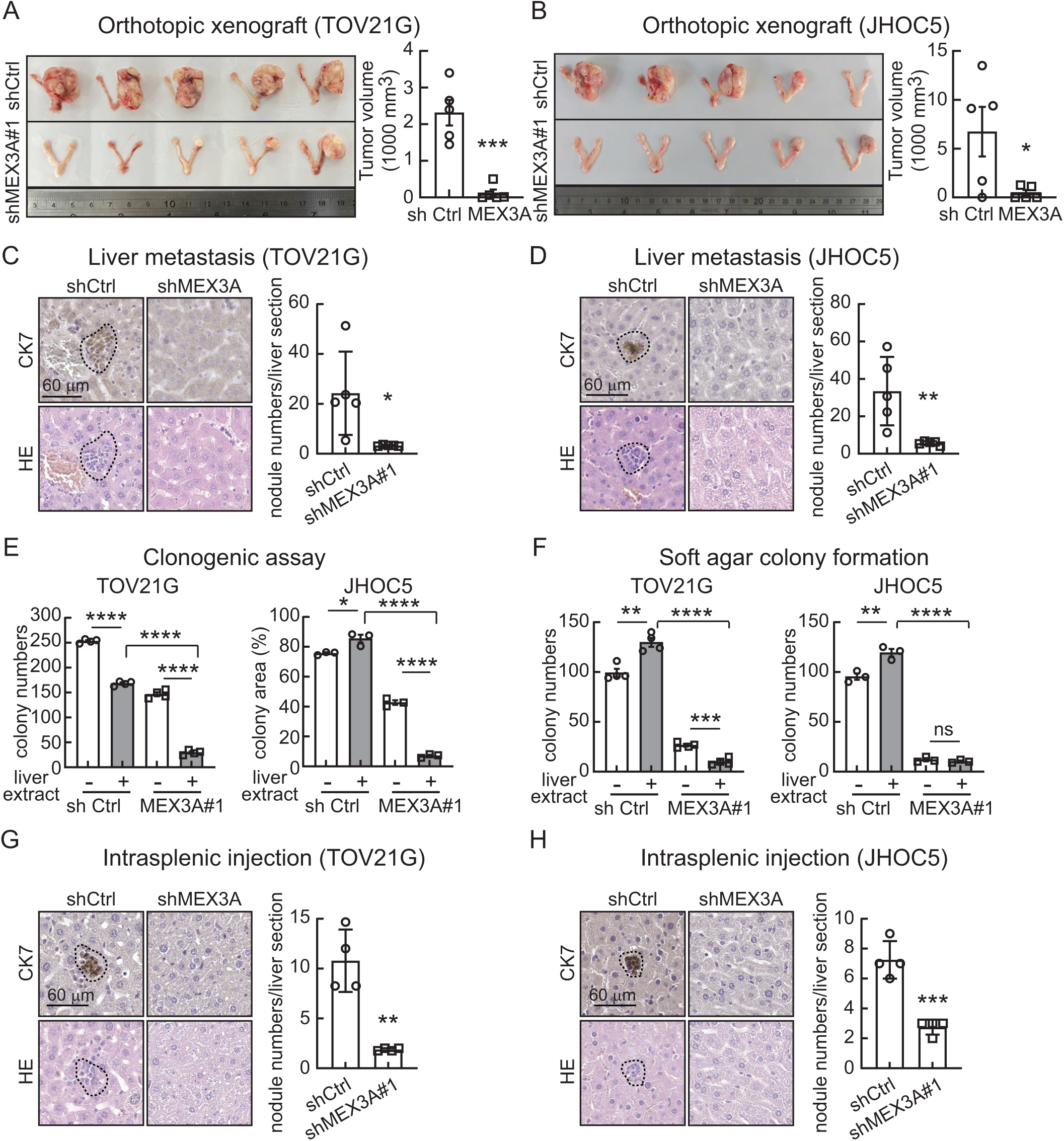
MEX3A promotes OCCC tumorigenesis and metastatic colonization in the liver. **(A)** Orthotopic xenograft models in NUDE mice using shCtrl or shMEX3A#1 TOV21G cells. Five mice were used for each group. Data are means ± SD, significant difference is based on unpaired *t* test of the tumor size 6 weeks after the *intra bursal* injection. ***, *P* < 0.001. **(B)** Orthotopic xenograft models in NUDE mice using shCtrl or shMEX3A#1 JHOC5 cells. Five mice were used for each group. Data are means ± SD, significant difference is based on unpaired *t* test of the tumor size 10 weeks after the *intra bursal* injection. *, *P* < 0.05. **(C)** Representative IHC images of liver sections from mice orthotopically injected with shCtrl or shMEX3A#1 TOV21G cells and quantification of liver nodule numbers in shCtrl and shMEX3A groups. Error bars indicate SD; *P* value was determined by unpaired *t* test. *, *P* < 0.05. **(D)** Representative IHC images of liver sections from mice orthotopically injected with shCtrl or shMEX3A#1 JHOC5 cells and quantification of liver nodule numbers in shCtrl and shMEX3A groups. Data formatting is as described for (C). **, *P* < 0.01. **(E)** Clonogenic assays using shCtrl or shMEX3A TOV21G and JHOC5 cells treated without or with liver extract. Data are shown as mean ± SD with *P* value based on unpaired *t* test (n = 4). *, *P* < 0.05; ****, *P* < 0.0001. The experiments were repeated 3 times. **(F)** Soft agar colony formation assays using shCtrl or shMEX3A TOV21G and JHOC5 cells treated without or with liver extract. Data are shown as mean ± SD with *P* value based on unpaired *t* test (n = 4). **, *P* < 0.01; ***, *P* < 0.001; ****, *P* < 0.0001; ns, not significant. The experiments were repeated 3 times. **(G)** Intrasplenic xenograft models in NUDE mice using shCtrl or shMEX3A#1 TOV21G cells. Four mice were used for each group. Data are means ± SD, significant difference is based on unpaired *t* test of the nodule numbers 3 weeks after the injection. **, *P* < 0.01. **(H)** Intrasplenic xenograft models in NUDE mice using shCtrl or shMEX3A#1 JHOC5 cells. Data formatting is as described for (G). ***, *P* < 0.001.

The liver constantly processes harmful substances by metabolism and detoxification, which often generates high level of toxic metabolites and reactive oxygen species (ROS). Such a stressful environment can lead to cell death or can inhibit cellular proliferation and tumorigenesis. To determine whether MEX3A contributed to tolerance of the liver microenvironment, clonogenic assays were performed using complete growth medium or medium containing liver extract from NUDE mice. Consistent with the xenograft assays, MEX3A-depleted cells had reduced rates of propagation and colony formation in liver extract media (Fig. 1E). Approximately 65% of the MEX3A-expressing TOV21G cells were able to form colonies while less than 20% of the MEX3A-depleted TOV21G cells were able to propagate in the liver extract containing medium (Fig. 1E). Similarly, MEX3A-expressing JHOC5 cells were able to survive in the liver extract containing medium while MEX3A-depleted JHOC5 cells were more sensitive to the liver extract treatment (Fig. 1E). We also evaluated the impact of liver extract on anchorage independent growth ability of OCCC cells using soft agar colony formation assays. As reported previously (26), MEX3A depletion significantly decreased soft agar colony forming ability of OCCC cells and liver extract further reduced colony numbers in MEX3A-depleted cells (Fig. 1F). Interestingly, in MEX3A-expressing cells, liver extract had the opposite effect as it led to an increase of anchorage-independent growth ability (Fig. 1F).

We then performed *intrasplenic* injection to bypass the initial steps of metastasis and thereby more directly evaluate whether MEX3A was required to form tumor nodules in the liver. Consistent with the data described above, MEX3A-depleted cells had reduced ability to form liver nodules compared to MEX3A-expressing cells (Fig. 1G and 1H). Together, these results suggested that MEX3A was not only required for tumor growth at the primary site, but was also critical for OCCC cells to survive and propagate in the stressful liver environment.

### MEX3A depletion leads to mitochondrial fragmentation and dysfunction in OCCC cells

Mitochondria fitness is critical for cancer progression. During metastasis, cancer cells need to precisely control mitochondrial dynamics to respond to changing energy demands and survive in different microenvironments. We found high levels of mitochondrial fragmentation in MEX3A-depleted cells. First, we observed that the length and width of mitochondria were significantly decreased (Fig. 2A-2D) while total mitochondrial mass was increased (Fig. 2E-2H) after MEX3A depletion in both TOV21G and JHOC5 cells. These data were indicative of increased mitochondria fission upon MEX3A depletion. We then examined the level of mitochondrial fission factor (MFF) (43), and its interacting GTPase, dynamin-related protein 1 (DRP1) (44), two essential factors that regulate mitochondrial fission. MEX3A depletion significantly increased MFF expression and increased DRP1 phosphorylation at Ser616, which is known to promote its mitochondrial localization for subsequent fission activity (Fig. 3). The elevation of these markers confirmed that MEX3A depletion led to increased mitochondrial fission.

**Figure 2.**
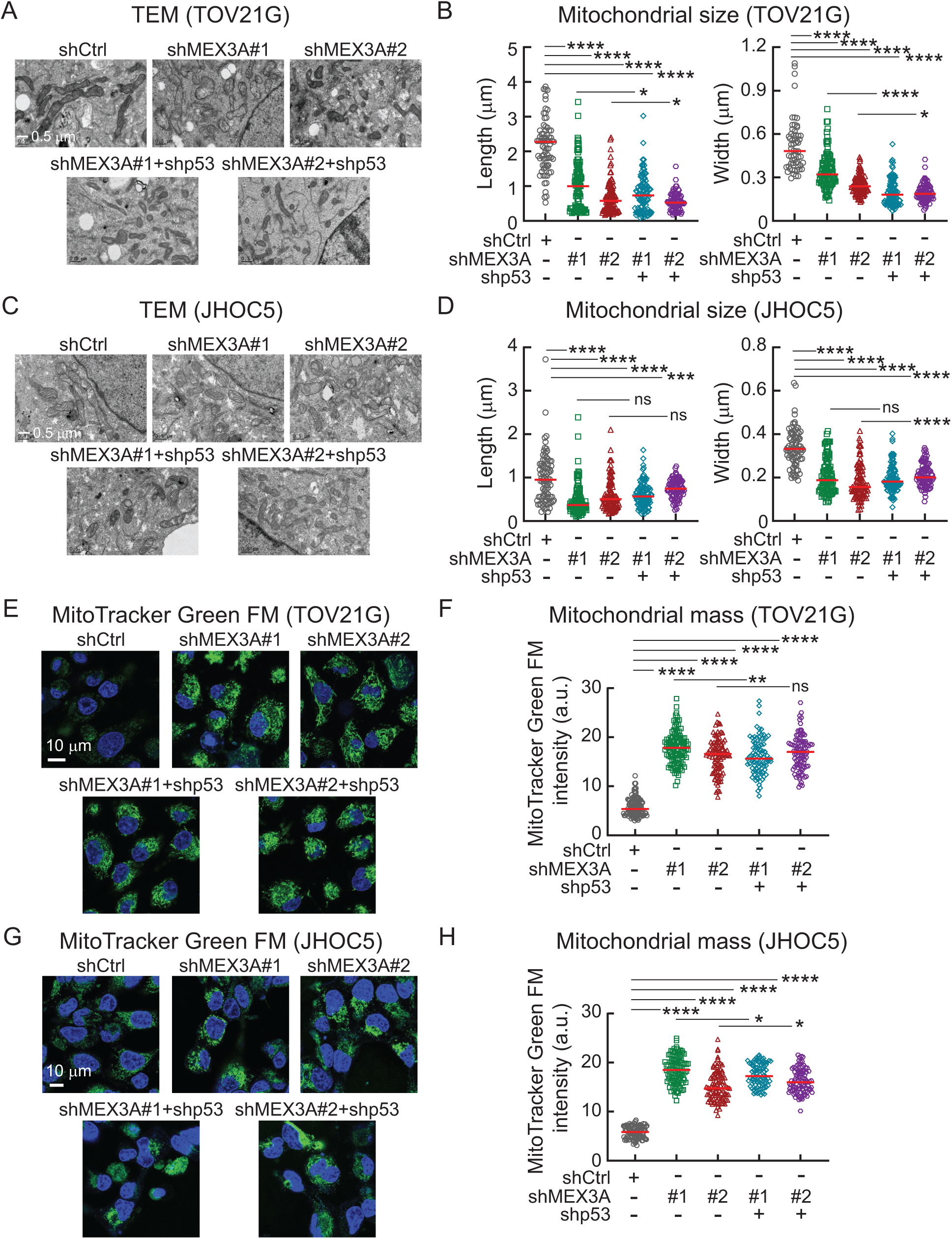
MEX3A depletion results in mitochondrial fragmentation in OCCC cells. **(A)** Representative TEM images of shCtrl, shMEX3A (#1 and #2), and MEX3A/p53 double knockdown (shMEX3A+shp53) TOV21G cells. (Scale bar, 0.5 µm.) **(B)** Quantification of mitochondrial length (Left) and width (Right) in shCtrl, shMEX3A, and shMEX3A+shp53 TOV21G cells. About 60 to 100 mitochondria were measured in each group. *P* value was determined by two-way ANOVA. *, *P* < 0.05; ****, *P* < 0.0001. **(C)** Representative TEM images of shCtrl, shMEX3A (#1 and #2), and MEX3A/p53 double knockdown (shMEX3A+shp53) JHOC5 cells. (Scale bar, 0.5 µm.) **(D)** Quantification of mitochondrial length (Left) and width (Right) in shCtrl, shMEX3A, and shMEX3A+shp53 JHOC5 cells. Data formatting is as described for (B). ***, *P* < 0.001; ****, *P* < 0.0001; ns, not significant. **(E)** Representative MitoTracker Green FM staining of shCtrl, shMEX3A (#1 and #2), and shMEX3A+shp53 TOV21G cells. (Scale bar, 10 µm.) **(F)** Quantification of MitoTracker Green FM intensity in shCtrl, shMEX3A, and shMEX3A+shp53 TOV21G cells. About 80-100 cells were measured in each group. *P* value was determined by two-way ANOVA. **, *P* < 0.01; ****, *P* < 0.0001; ns, not significant. **(G)** Representative MitoTracker Green FM staining of shCtrl, shMEX3A (#1 and #2), and shMEX3A+shp53 JHOC5 cells. (Scale bar, 10 µm.) **(H)** Quantification of MitoTracker Green FM intensity in shCtrl, shMEX3A, and shMEX3A+shp53 JHOC5 cells. Data formatting is as described for (F). *, *P* < 0.05; ****, *P* < 0.0001.

**Figure 3.**
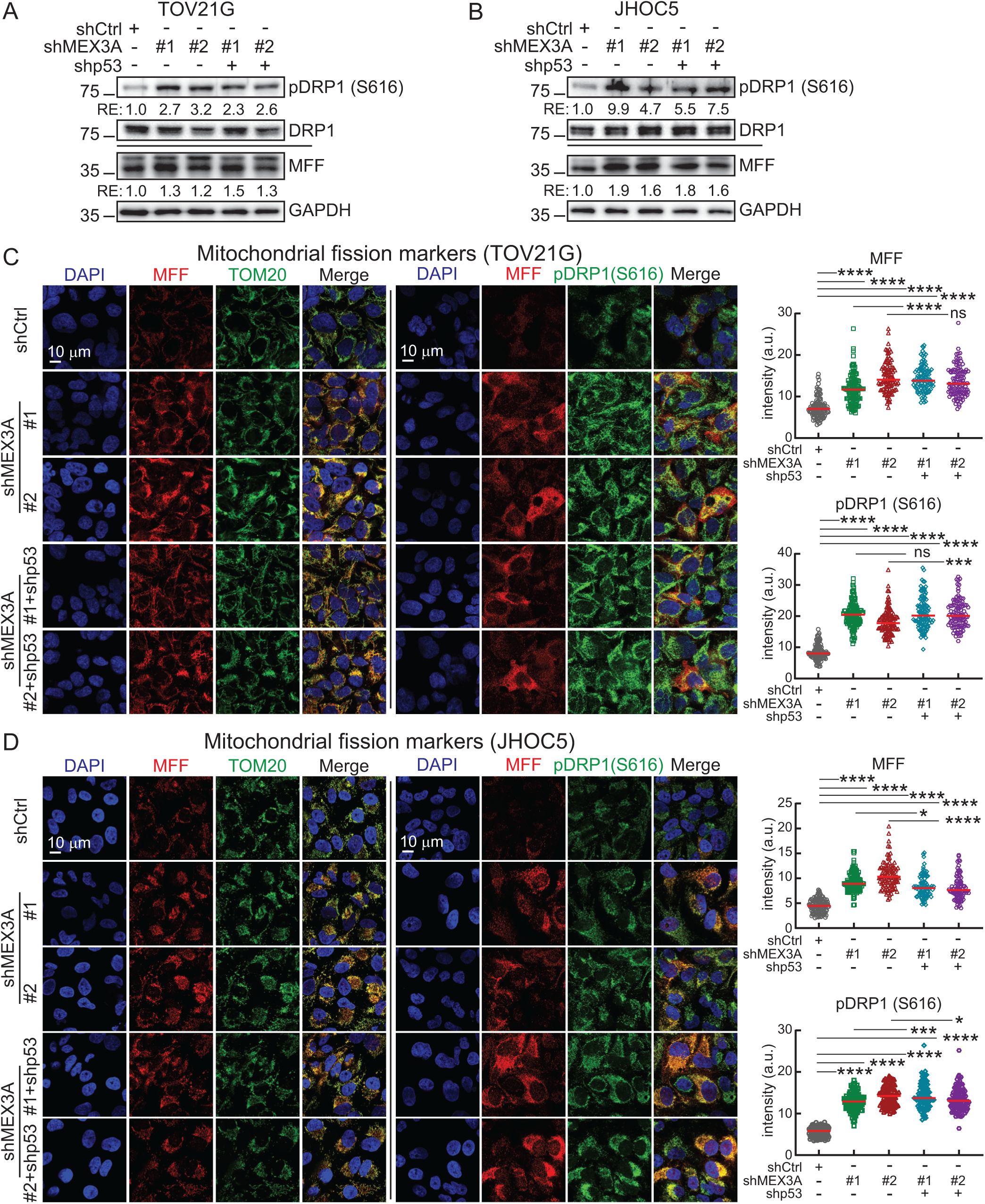
MEX3A depletion increases mitochondrial fission in OCCC cells. **(A)** Immunoblot of pDRP1 (S616), DRP1, and MFF in shCtrl, shMEX3A and shMEX3A+shp53 TOV21G cells. GAPDH was used as a loading control. Blots shown are from one representative experiment of three replicates. RE, relative expression level. **(B)** Immunoblot of pDRP1 (S616), DRP1, and MFF in shCtrl, shMEX3A and shMEX3A+shp53 JHOC5 cells. GAPDH was used as a loading control. Blots shown are from one representative experiment of three replicates. RE, relative expression level. **(C)** Representative immunofluorescence (IF) images of DAPI, MFF, TOM20 and pDRP1 (S616), and quantification of MFF and pDRP1 (S616) intensity in shCtrl, shMEX3A and shMEX3A+shp53 TOV21G cells. About 90-100 cells were measured in each group. *P* value was determined by two-way ANOVA. ****, *P* < 0.0001; ns, not significant. (Scale bar, 10 µm.) **(D)** Representative IF images of DAPI, MFF, TOM20 and pDRP1 (S616), and quantification of MFF and pDRP1 (S616) intensity in shCtrl, shMEX3A and MEX3A+shp53 JHOC5 cells. Data formatting is as described for (C). *, *P* < 0.05; ***, *P* < 0.001; ****, *P* < 0.0001.

Since p53 has also been shown to affect mitochondrial fission (45,46) and p53 protein level increases upon MEX3A depletion (Fig. S1E) (26), we generated MEX3A/p53 double knockdown cells (Fig. S1E) to test whether the fission phenotype observed in MEX3A depleted cells was due to p53 upregulation. The results showed that p53 knockdown did not fully rescue the MEX3A-depletion mediated mitochondrial fission phenotypes including mitochondrial morphology (Fig. 2A-2D), mass (Fig. 2E-2H) and fission markers (Fig. 3). Thus, the function of MEX3A in mitochondrial dynamics was largely independent of p53.

Mitochondrial integrity, maintained by dynamic fusion and fission, is critical for cellular homeostasis. However, chronic mitochondrial fragmentation is often associated with decreased mitochondrial function and thus is detrimental to cellular function (47). Consistent with this, several assays indicated that the fragmented mitochondria in MEX3A-depleted OCCC cells became dysfunctional. Using JC-1 as a mitochondrial membrane potential indicator, we found that membrane potential was decreased significantly in MEX3A-depleted cells (Fig. 4A and 4B). In addition, these cells had increased superoxide level (Fig. 4C and 4D), decreased NAD^+^/NADH ratio (Fig. 4E and 4F), impaired OXPHOS (Fig. 4G and 4H), and reduced ATP production (Fig. 4I and 4J), confirming that MEX3A depletion resulted in defective mitochondria. These changes in mitochondrial fitness persisted in MEX3A-depleted cells when p53 was knocked down, indicating that these effects were independent of p53. It is worth noting that the link between MEX3A depletion, mitochondrial fission and impaired OXPHOS was observed in OCCC but not in HGSOC (Fig. S2), indicating a subtype specific effect.

**Figure 4.**
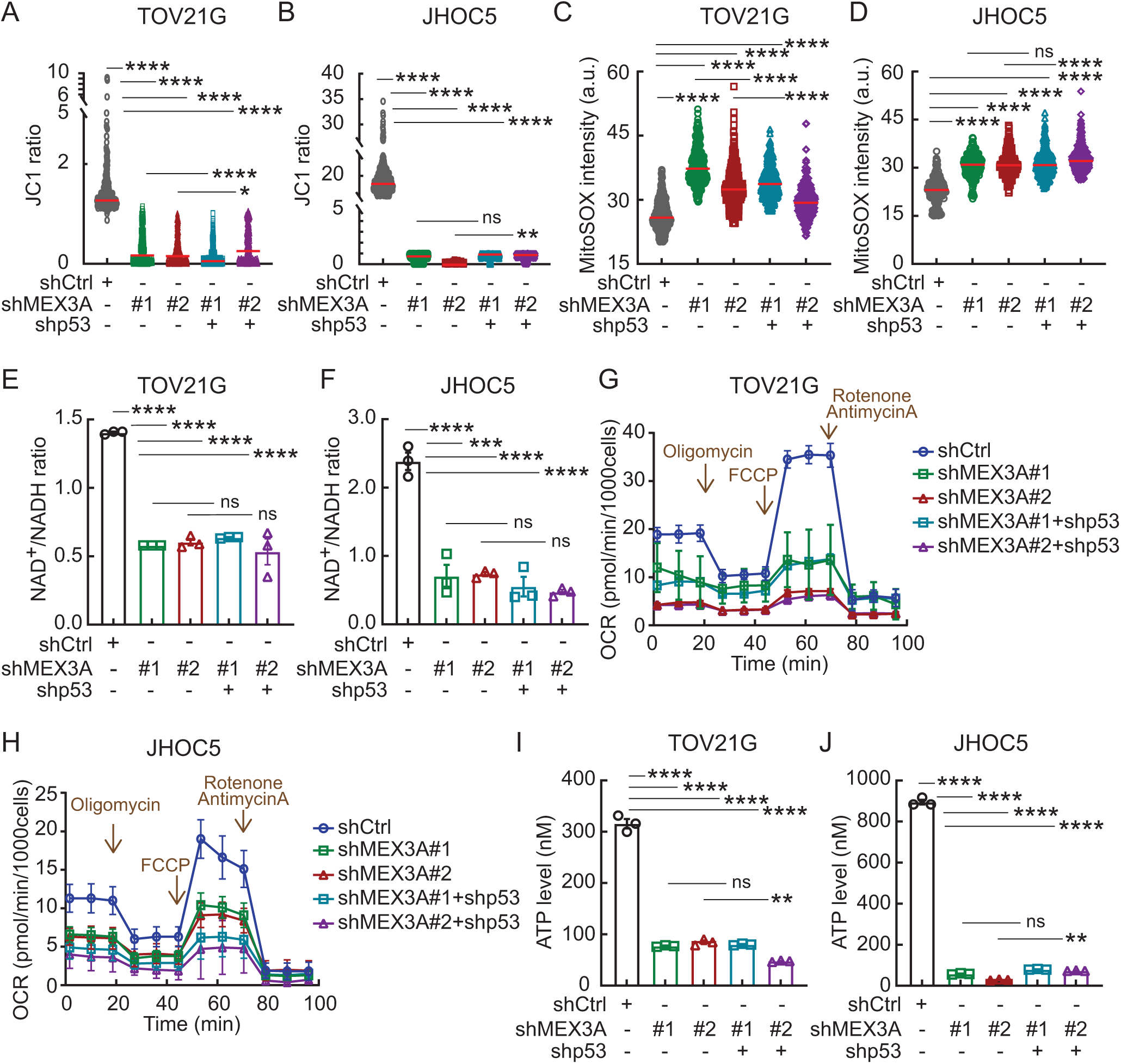
MEX3A depletion impairs mitochondrial function in OCCC cells. **(A)** Mitochondrial membrane potential indicated by JC-1 ratio in shCtrl, shMEX3A, and shMEX3A+shp53 TOV21G cells. More than 250 cells were measured in each group. *P* value was determined by two-way ANOVA. *, *P* < 0.05; ****, *P* < 0.0001. **(B)** JC-1 ratio in shCtrl, shMEX3A, and shMEX3A+shp53 JHOC5 cells. Data formatting is as described for (A). ****, *P* < 0.0001. **(C)** Superoxide level indicated by MitoSOX intensity in shCtrl, shMEX3A, and shMEX3A+shp53 TOV21G cells. More than 250 cells were measured in each group. *P* value was determined by two-way ANOVA. ****, *P* < 0.0001. **(D)** MitoSOX intensity in shCtrl, shMEX3A, and shMEX3A+shp53 JHOC5 cells. Data formatting is as described for (C). ****, *P* < 0.0001; ns, not significant. **(E)** NAD^+^/NADH ratio in shCtrl, shMEX3A, and shMEX3A+shp53 TOV21G cells. *P* value was determined by two-way ANOVA. ****, *P* < 0.0001; ns, not significant. **(F)** NAD^+^/NADH ratio in shCtrl, shMEX3A, and shMEX3A+shp53 JHOC5 cells. Data formatting is as described for (E). ***, *P* < 0.001; ****, *P* < 0.0001; ns, not significant. **(G)** OXPHOS activity in shCtrl, shMEX3A, and shMEX3A+shp53 TOV21G cells. Graph shown is from one representative experiment of three replicates. **(H)** OXPHOS activity in shCtrl, shMEX3A, and shMEX3A+shp53 JHOC5 cells. Graph shown is from one representative experiment of three replicates. **(I)** ATP level in shCtrl, shMEX3A, and shMEX3A+shp53 TOV21G cells. Data are shown as mean ± SD with *P* value based on two-way ANOVA (n = 3). **, *P* < 0.01; ****, *P* < 0.0001; ns, not significant. The experiments were repeated 3 times. **(J)** ATP level in shCtrl, shMEX3A, and shMEX3A+shp53 JHOC5 cells. Data formatting is as described for (I). **, *P* < 0.01; ****, *P* < 0.0001; ns, not significant.

### MEX3A contributes to ETC supercomplex assembly

RNA-sequencing (RNA-seq) analysis using OCCC cells with or without MEX3A depletion was performed to identify transcriptome effects of MEX3A disruption. Gene Set Enrichment Analysis (GSEA) showed that more than 30% of the OXPHOS genes in the cancer hallmark was significantly correlated with MEX3A expression in TOV21G cells (Fig.5A). Differential expression analysis found that these OXPHOS genes, including genes encoding mitochondrial electron transport chain (ETC) complex I core subunits, such as NDUFS1 and NDUFS2, and accessory subunits, such as NDUFB7 and NDUFC2, were downregulated in MEX3A-depleted cells (Fig. 5B and 5C). These data indicated that MEX3A depletion may lead to a decreased level of complex I subunits and therefore reduced assembly of functional complex I. It has been reported that structural and functional defects in one ETC complex can trigger dysfunction or instability of other complexes, and disrupt the formation, structural and functional integrity of supercomplexes (48). Consistent with this, Blue Native polyacrylamide gel electrophoresis (BN-PAGE) showed that the amount of ETC complexes were substantially reduced in MEX3A-depleted cells, particularly complex I+III+IV, I and III+IV/V (Fig. 5D and 5E). Together with the reduced OXPHOS level and other mitochondrial functions observed in the MEX3A-depleted cells (Fig. 4), these results indicate that MEX3A control of mitochondrial fitness occurs, in part, via control of ETC complex I subunit expression.

**Figure 5.**
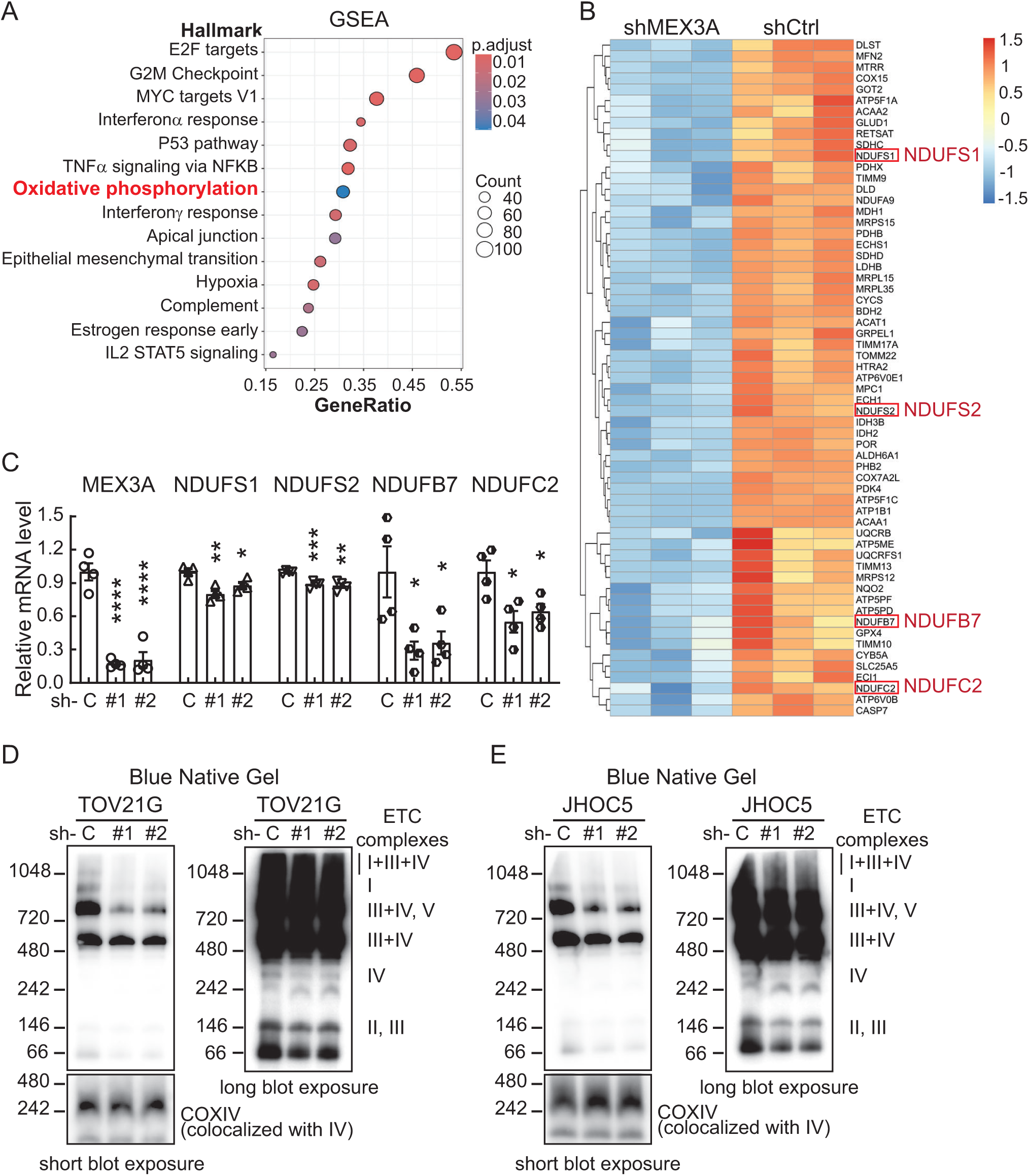
MEX3A regulates mitochondria-related genes and is critical for ETC supercomplex assembly. **(A)** Gene Set Enrichment Analysis (GSEA) of downregulated genes in MEX3A-depleted TOV21G cells. The OXPHOS pathway was found in the top 14 enriched pathways in the MEX3A-depleted group. **(B)** Heatmap of OXPHOS related genes in TOV21G cells without or with MEX3A depletion. **(C)** qRT-PCR analysis of NDUFS1, NDUFS2, NDUFB7 and NDUFC2 expression in shCtrl (C) or shMEX3A (#1, #2) JHOC5 cells. Cyclophilin was used as an internal control. Three independent experiments were performed and data are means ± SD from one representative experiment (n =4). Significant differences are based on unpaired *t* test. *, *P* < 0.05; **, *P* < 0.01; ***, *P* < 0.001; ****, *P* < 0.0001. **(D)** Blue-Native PAGE of respiratory super complexes in shCtrl and shMEX3A (#1 and #2) TOV21G cells. Blots shown are from one representative experiment of three replicates. **(E)** Blue-Native PAGE of respiratory super complexes in shCtrl and shMEX3A (#1 and #2) JHOC5 cells. Blots shown are from one representative experiment of three replicates.

### MEX3A expression allows OCCC cells to recover from mitophagy and survive in stressful microenvironments

During tumor progression and metastasis, cancer cells need to cope with different stressors, including hypoxia, nutrient deprivation, low pH, and high ROS, all of which can damage mitochondria. To maintain proper energy metabolism and growth in different microenvironments, cancer cells rely on mitophagy to remove damaged or excessive mitochondria. Importantly, after mitochondrial removal, cells must produce new and healthy mitochondria. Otherwise, mitochondrial depletion will lead to insufficient energy production and subsequent cell death (49). In addition to its critical role in maintenance of mitochondrial function, MEX3A effect on mitophagy recovery could also be a reason that it is essential for OCCC liver metastasis. As cellular stress responses can occur rapidly, we first performed a three-day time course cell proliferation assay to confirm that MEX3A depletion made the cells hypersensitive to liver extract (Fig. 6A and S3A). While depletion of MEX3A itself decreased cell survival, consistent with previous report (26), the addition of liver extract increased cell mortality to nearly 90% for MEX3A-depleted TOV21G cells (Fig. 6A) and 80% for MEX3A-depleted JHOC5 cells (Fig. S3A) but had no effect on control cells. These results, together with the liver extract inhibition of clonogenic and anchorage-independent growth ability in MEX3A-depleted cells (Fig. 1), indicated that MEX3A was the key factor which allowed cells to cope with stress factors present in the liver. Liver extract induced mitophagy and autophagy in MEX3A-expressing OCCC cells, as indicated by an increase in the key mitophagy markers PTEN-induced kinase 1 (PINK1), phosphorylated PINK1 (p-PINK1), PARKIN, and p62 (also known as SQSTM1), as well as the autophagy marker LC3 (Microtubule-associated protein light chain 3) within 4 hours after treatment (Fig. 6B and S3B). The basal level of these stress markers was higher in MEX3A-depleted cells (Fig. 6C and S3C), likely due to accumulation of fragmented mitochondria (Fig. 2); however, addition of liver extract clearly induced further autophagy in MEX3A-depleted cells as indicated by the increased level of LC3 (Fig. 6B). Notably, in MEX3A-depleted cells, mitophagy markers were significantly decreased and mitochondrial marker TOM20 was severely depleted 12 hours after treatment (Fig. 6B and S3B). By 24 hours, mitochondrial mass was significantly decreased in these cells (Fig. 6D and S3D), suggesting that generation of new mitochondria was inhibited. In contrast. such reduction of mitochondrial mass was not observed in the MEX3A-expressing cells (Fig. 6D and S3D). Since our data indicated that MEX3A has a previously unsuspected role in mitochondrial fitness (Fig. 2-5) and is required for OCCC liver metastasis (Fig. 1), we hypothesize that MEX3A-expressing cells can tolerate and recover from stress-induced mitophagy; while MEX3A-depleted cells cannot.

**Figure 6.**
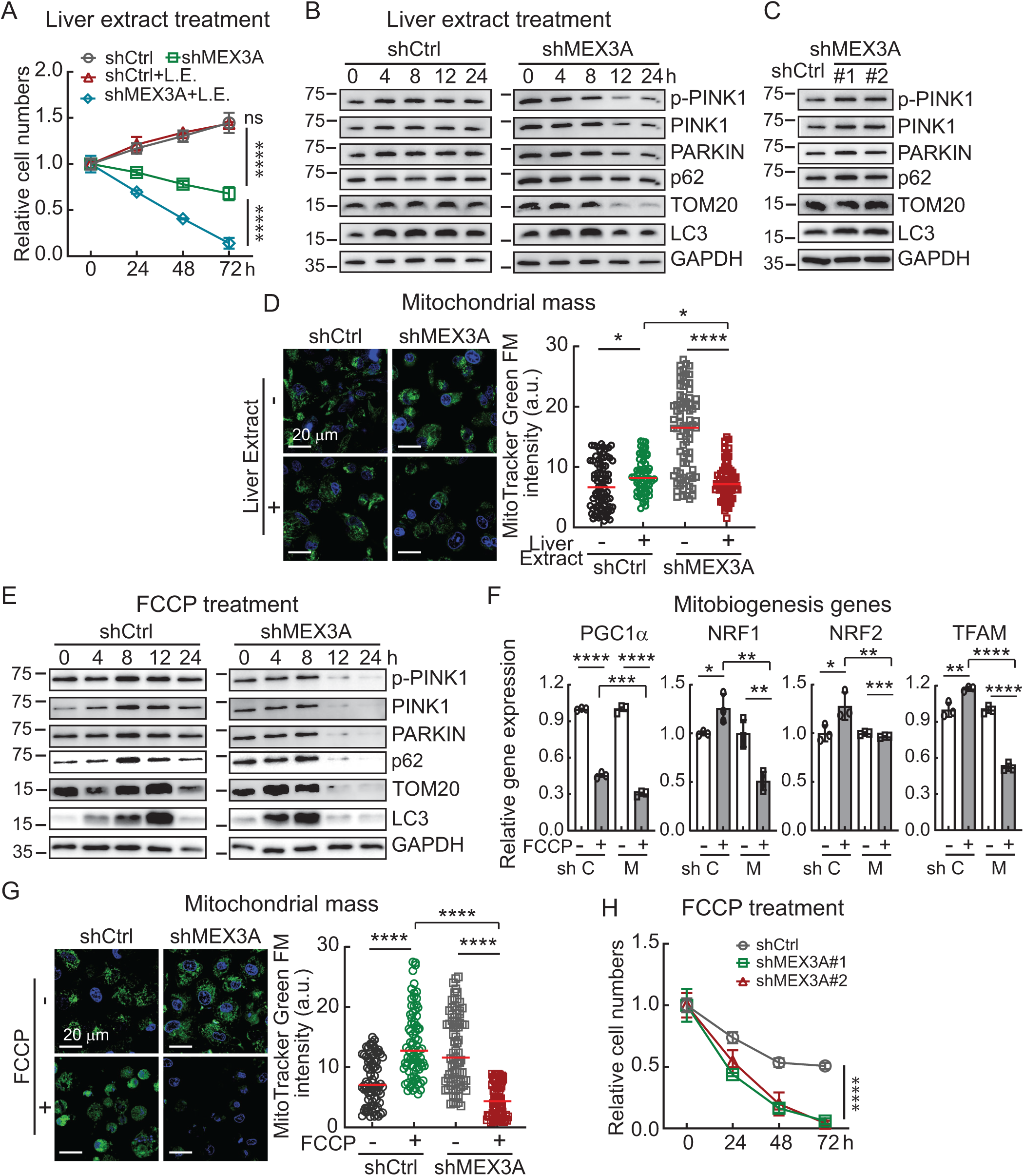
MEX3A is required for OCCC cells to recover from mitophagy and survive stress from liver extract or FCCP treatment. **(A)** Cell survival time course assay using liver extract (L.E.) treated shCtrl or shMEX3A TOV21G cells. Cell numbers were counted every day for 3 days. Data are shown as mean ± SD with *P* value based on two-way ANOVA-test (n = 3). ****, *P* < 0.0001; ns, not significant. The experiments were repeated 3 times. **(B)** Time course assay using liver extract (10%) treated shCtrl or shMEX3A TOV21G cells. The levels of PINK1, pPINK1, PARKIN, p62, LC3 and TOM20 were determined using IB analysis. GAPDH was used as a loading control. **(C)** Immunoblot of PINK1, pPINK1, PARKIN, p62, and LC3 and TOM20 in shCtrl and shMEX3A TOV21G cells. GAPDH was used as a loading control. Blots shown are from one representative experiment of three replicates. RE, relative expression level. **(D)** Representative MitoTracker Green FM staining of shCtrl, shMEX3A (#1 and #2) TOV21G cells treated with liver extract (10%). (Scale bar, 20 µm.) Quantification of MitoTracker Green FM intensity in shCtrl and shMEX3A TOV21G cells. About 80-100 cells were measured in each group. *P* value was determined by unpaired *t* test. *, *P* < 0.05; ****, *P* < 0.0001. **(E)** Time course assay using FCCP (10 µM) treated shCtrl or shMEX3A TOV21G cells. Data formatting as described for (B). **(F)** qRT-PCR analysis of PGC1α, NRF1, NRF2, and TFAM expression in shCtrl or shMEX3A TOV21G cells treated with 10 µM FCCP. Cyclophilin was used as an internal control. Three independent experiments were performed and data are means ± SD from one representative experiment (n =3). Significant differences are based on unpaired *t* test. *, *P* < 0.05; **, *P* < 0.01; ***, *P* < 0.001; ****, *P* < 0.0001. **(G)** Representative MitoTracker Green FM staining of shCtrl, shMEX3A (#1 and #2) TOV21G cells treated with 10 µM FCCP. (Scale bar, 20 µm.) Quantification of MitoTracker Green FM intensity in shCtrl and shMEX3A TOV21G cells. About 80-100 cells were measured in each group. *P* value was determined by unpaired *t* test. ****, *P* < 0.0001. **(H)** Cell survival time course assay using 10 µM FCCP treated shCtrl or shMEX3A TOV21G cells. Cell numbers were counted every day for 3 days. Data are shown as mean ± SD with *P* value based on two-way ANOVA-test (n = 3). ****, *P* < 0.0001. The experiments were repeated 3 times.

Since there are several stress factors, such as inflammatory cytokines and metabolites, in the liver extract that could cause stresses other than mitophagy, we used FCCP to more specifically depolarize mitochondria and induce mitophagy in OCCC cells (Fig. 6E and S3E). In MEX3A-expressing TOV21G cells, levels of key mitophagy markers were elevated and peaked at 8 hours after FCCP treatment (Fig. 6E). LC3 was also increased and peaked at 12 hours after FCCP treatment (Fig. 6E). These markers all decreased to basal levels at 24 hours after mitophagy induction (Fig. 6E), indicating that the damaged mitochondria had been removed. The maintenance of TOM20 levels in MEX3A-expressing cells indicated that healthy, or newly generated, mitochondria continued to support cellular needs after mitophagy was completed. For TOV21G MEX3A-depleted cells, the rapid induction of LC3 indicated that mitophagy occurred (Fig. 6E). However, at later time points, MEX3A-depleted cells had greatly reduced levels of all mitochondrial proteins, including TOM20, indicative of an inability to generate new mitochondria (Fig. 6E). Similar patterns were observed in MEX3A-expressing versus MEX3A-depleted JHOC5 cells (Fig. S3E).

We also found that genes involved in mitochondrial biogenesis including NRF1, NRF2 and TFAM were upregulated in MEX3A-expressing cells 24 hours after FCCP treatment; but were significantly decreased in MEX3A-depleted cells (Fig. 6F and S3F). Although PGC1α was reduced in both cells after FCCP treatment, the level of PGC1α was much lower in MEX3A-depleted cells (Fig. 6F and S3F). In addition, a more severe decrease in mitochondrial mass was found in MEX3A-depleted cells after FCCP treatment (Fig. 6G and S3G). These results indicated that damaged mitochondria were removed in both MEX3A-expressing and MEX3A-depleted cells; however, MEX3A-depleted cells failed to generate new mitochondria. Consistent with this, MEX3A-depleted OCCC cells failed to survive FCCP treatment that produced only partial mortality in MEX3A-expressing cells (Fig. 6H and S3H). Together, our observations show that MEX3A is required for OCCC cells to recover from mitophagy, survive and propagate in the liver microenvironment.

## Discussion

Cancer progression requires sufficient energy production to sustain tumor growth and metastasis, as well as dynamic metabolic reprogramming to cope with the stresses induced by different microenvironments. For a long time, aerobic glycolysis (a.k.a. the Warburg effect) has been seen as a cancer hallmark and a special metabolic pathway for cancer to survive stressful conditions. For example, OCCC cells express high levels of HNF1β to enhance aerobic glycolysis and survive in the endometriotic cyst which contains high levels of free iron and ROS (50). Recently, evidence accumulated across many experiments shows that glycolysis and OXPHOS can occur concurrently to support cancer cell proliferation (51). Furthermore, when cancer cells become more malignant and chemoresistant, OXPHOS upregulation is correlated with poor outcome (51,52). In OC, invasive cells often have higher OXPHOS activity compared to less invasive OC cells or normal ovarian epithelial cells (53,54). In OCCC, upregulated expression of OXPHOS-related genes, elevated mitochondrial membrane potential and respiration, as well as increased mitochondrial number and increased cristae/outer membrane surface ratio have been observed, indicative of a high energy demand in OCCC cells (55). However, intrinsic factors responsible for promoting mitochondrial fitness to maintain OXPHOS, and the relationship between mitochondrial fitness and OCCC progression have remained largely unknown. Meanwhile, several lines of evidence indicate that aberrant expression of MEX3A upregulates multiple pathways to drive tumor initiation, progression and metastasis in many types of cancer (26,32,33). However, those prior studies did not link MEX3A to cancer metabolism. Prior to our study, there was only one report showing that MEX3A could promote colorectal cancer angiogenesis via activation of glycolysis (56). Our cell-based experiments and animal models demonstrate that MEX3A-mediated maintenance of mitochondrial fitness and OXPHOS is required for OCCC tumorigenesis and is especially critical for liver metastasis.

RBPs similar to MEX3A participate in both cytoplasmic and mitochondrial post-transcriptional regulation to fine-tune mitochondrial respiration in response to different environmental stressors and provide adequate energy for cells (57). Based on our transcriptomic analyses, MEX3A may have even further impacts on mitochondrial dynamics and metabolism in OCCC cells in addition to promoting Complex I subunit expression (Fig. 5B; Table S5). For example, MFN2, a mitochondrial outer membrane GTPase that mediates mitochondrial clustering and fusion, was downregulated in MEX3A-depleted cells. MFN2 deficiency results in a fragmented mitochondrial network and disruption of ER-mitochondria tethering which leads to mitochondrial dysfunction and elevated ER stress (58). Also, many genes critical for the TCA cycle and redox balance, such as isocitrate dehydrogenase (IDH2, IDH3B), pyruvate dehydrogenase X (PDHX), succinate dehydrogenase (SDHC), and glutathione peroxidase 4 (GPX4), were downregulated in MEX3A depleted cells. However, it remains unclear whether MEX3A regulates these genes directly via its RNA binding activity, or indirectly modulates their upstream regulators through its E3 ligase function. Moreover, MEX3A effects on mitochondria were observed in OCCC cells but not in HGSOC cells (Fig. S2), suggesting that MEX3A oncogenic function is context dependent. Identification of OCCC-specific factors that cooperate with MEX3A to regulate mitochondrial fitness is an important topic for further investigation. Also, it will be of interest to determine whether the role of MEX3A-mitochondrial regulation identified in our study is also important for other cancer types that rely on OXPHOS for metastasis.

In summary, our study demonstrates that MEX3A maintains mitochondrial fitness to support high levels of OXPHOS needed for OCCC tumor growth and liver metastasis. Since OXPHOS inhibitors have low selectivity between normal and cancer cells and can cause potent and systemic effects in patients (59), our findings indicate that it may instead be promising to target MEX3A as a way to selectively inhibit mitochondrial function in cancer cells. Together with our previous demonstration that MEX3A suppressed p53 in OCCC (26), results presented here demonstrate that OCCC dependence on MEX3A for both tumor growth and liver metastasis is a critical vulnerability that can be used to develop new OCCC treatment strategies.

## Acknowledgments

This work was supported by the Taiwan National Science and Technology Council [NSTC 111-2314-B-001-012], [NSTC 112-2314-B-001-006] and [NSTC 113-2314-B-001-012] to Wendy W. Hwang-Verslues. The authors would like to thank the following core facilities at Academia Sinica: the National RNAi Core Facility for providing shRNA reagents and related services; the Bioinformatics Core and Imaging Core at the Institute of Molecular Biology for providing the RNA-seq analysis and TEM services, respectively; the DNA Sequencing Core Facility of the Institute of Biomedical Sciences [funded by Academia Sinica Core Facility and Innovative Instrument Project (AS-CFII-113-A12)] for DNA sequencing analysis; the Biochemistry Core Facility at the Institute of Cellular and Organismic Biology for Seahorse Analyzer service; the Mass Spectrometry Core Facility and Histology Core Facility at Genomics Research Center for technical support; the Advanced Optics Microscope Core Facility (funded by AS-CFII-114-A3) for microscope imaging technical support, and Academia Sinica SPF Animal Facility (AS-CFII-113-A7) for providing animal support.

