## Supplementary material for "MEX3A control of mitochondrial fitness is essential for ovarian clear cell carcinoma tumorigenesis and liver metastasis": Figure S1-S3, Table S1-S5

**This PDF file includes:**

Figures S1 to S3
Tables S1 to S3
Table S4 and S5 are in separate xlsx files.

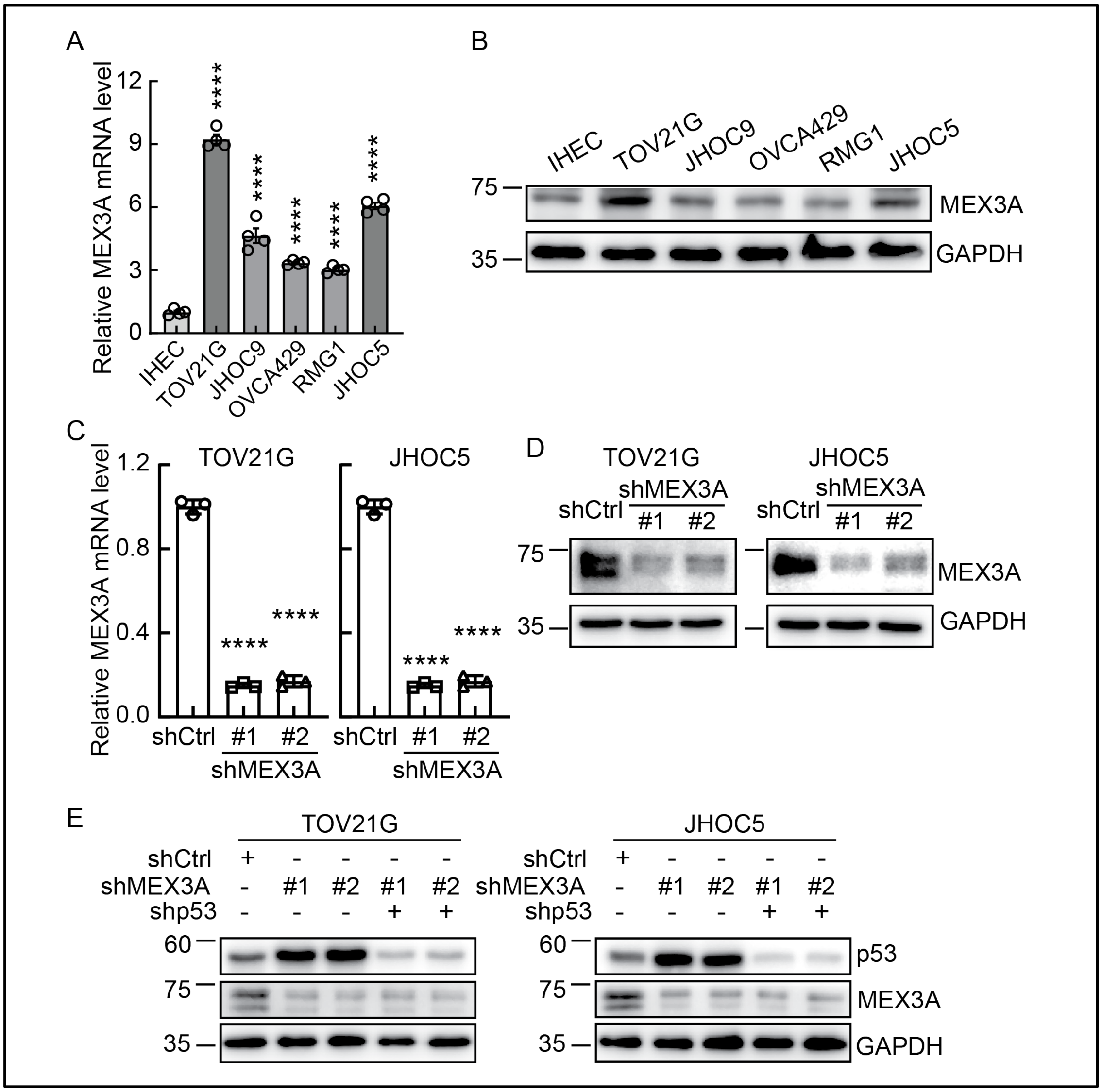

**Figure S1.** MEX3A expression in OCCC cell lines.

**(A)** qRT-PCR analysis of MEX3A level in IHEC and OCCC cells. Cyclophilin was used as an internal control. Three independent experiments were performed and data are means ± SD from one representative experiment (n = 4). ****, $P < 0.0001$. Significant differences are based on unpaired *t* test.

(**B**) IB of MEX3A protein expression with GAPDH as a loading control. Blots shown are from one representative experiment of three replicates.

**(C)** qRT-PCR analysis of MEX3A mRNA level in TOV21G and JHOC5 cells without (shCtrl) or with MEX3A depletion (shMEX3A#1 or shMEX3A#2). Cyclophilin was used as an internal control. Three independent experiments were performed and data are means ± SD from one representative experiment (n =3). ****, $P < 0.0001$. Significant differences are based on unpaired *t* test.

**(D)** IB of MEX3A protein level in TOV21G and JHOC5 cells without (shCtrl) or with MEX3A depletion (shMEX3A#1 or shMEX3A#2). GAPDH was used as a loading control. Blots shown are from one representative experiment of three replicates.

**(E)** IB of MEX3A and p53 protein level in TOV21G and JHOC5 cells without (shCtrl) or with MEX3A and p53 depletion (shMEX3A#1/shp53 or shMEX3A#2/shp53). GAPDH was used as a loading control. Blots shown are from one representative experiment of three replicates.

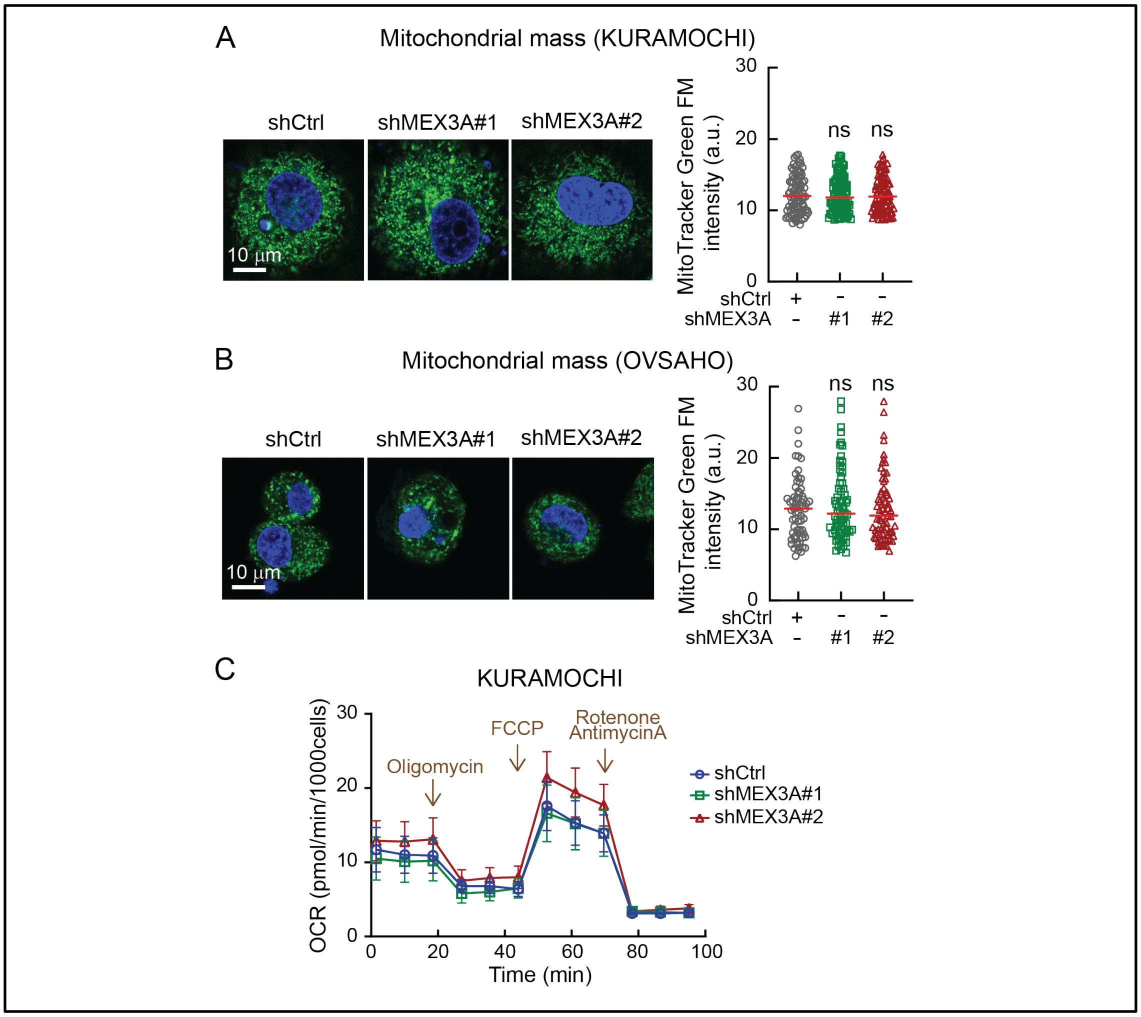

**Figure S2.** MEX3A depletion does not change mitochondrial mass or impair mitochondrial function in HGSOC cell lines.

**(A)** Representative MitoTracker Green FM staining of shCtrl and shMEX3A (#1 and #2) KURAMOCHI cells. (Scale bar, 10 µm.) Quantification of MitoTracker Green FM intensity in shCtrl and shMEX3A KURAMOCHI cells. About 90-100 cells were measured in each group. *P* value was determined by unpaired *t* test. ns, not significant.

**(B)** Representative MitoTracker Green FM staining of shCtrl and shMEX3A (#1 and #2) OVSAHO cells. (Scale bar, 10 µm.) Quantification of MitoTracker Green FM intensity in shCtrl and shMEX3A OVSAHO cells. About 70-80 cells were measured in each group. *P* value was determined by unpaired *t* test. ns, not significant.

**(C)** OXPHOS activity in shCtrl and shMEX3A KURAMOCHI cells. Graph shown is from one representative experiment of three replicates.

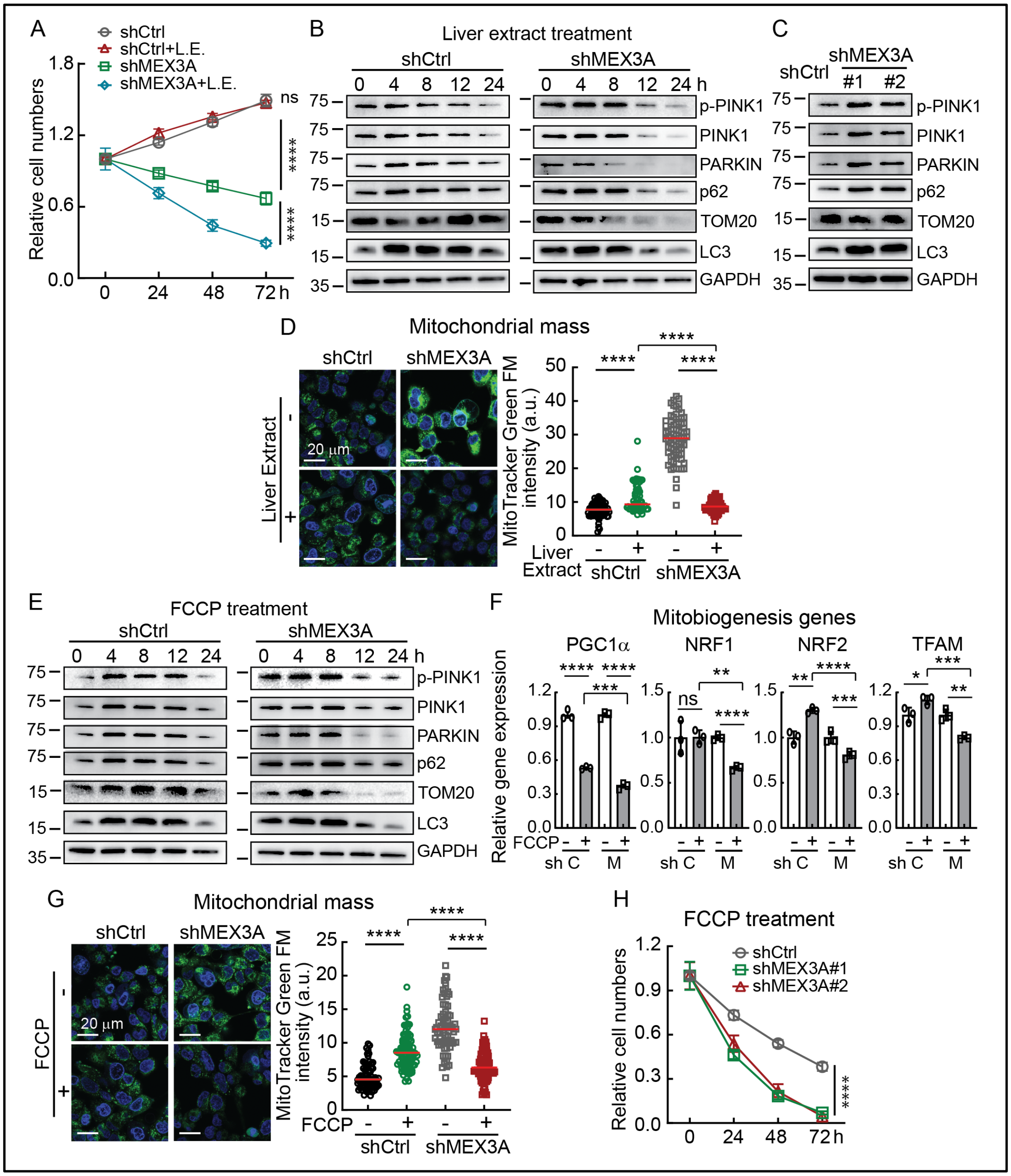

**Figure S3.** MEX3A is required for OCCC cells to recover from mitophagy.

**(A)** Cell survival time course assay using liver extract (10%) treated shCtrl or shMEX3A JHOC5 cells. Cell numbers were counted every day for 3 days. Data are shown as mean

± SD with *P* value based on two-way ANOVA-test (n = 3). ****, P < 0.0001. The experiments were repeated 3 times.

**(B)** Time course assay using liver extract (10%) treated shCtrl or shMEX3A JHOC5 cells. The levels of PINK1, pPINK1, PARKIN, p62, LC3 and TOM20 were determined using IB analysis. GAPDH was used as a loading control.

**(C)** Immunoblot of PINK1, pPINK1, PARKIN, p62, and LC3 and TOM20 in shCtrl and shMEX3A JHOC5 cells. GAPDH was used as a loading control. Blots shown are from one representative experiment of three replicates.

**(D)** Representative MitoTracker Green FM staining of shCtrl, shMEX3A (#1 and #2) JHOC5 cells treated with liver extract (10%). (Scale bar, 20 μm.) Quantification of MitoTracker Green FM intensity in shCtrl and shMEX3A JHOC5 cells. About 50-80 cells were measured in each group. *P* value was determined by unpaired *t* test. ****, *P* < 0.0001.

**(E)** Time course assay using FCCP (10 μM) treated shCtrl or shMEX3A JHOC5 cells. Data formatting as described for (A).

**(F)** qRT-PCR analysis of PGC1α, NRF1, NRF2, and TFAM expression in shCtrl or shMEX3A JHOC5 cells treated with 10 μM FCCP. Cyclophilin was used as an internal control. Three independent experiments were performed and data are means ± SD from one representative experiment (n =3). Significant differences are based on unpaired *t* test. *, *P* < 0.05; **, *P* < 0.01; ***, *P* < 0.001; ****, *P* < 0.0001; ns, not significant.

**(G)** Representative MitoTracker Green FM staining of shCtrl, shMEX3A (#1 and #2) JHOC5 cells treated with 10 μM FCCP. (Scale bar, 20 μm.) Quantification of MitoTracker Green FM intensity in shCtrl and shMEX3A JHOC5 cells. About 80-100 cells were measured in each group. *P* value was determined by unpaired *t* test. ****, *P* < 0.0001.

**(H)** Cell survival time course assay using 10 μM FCCP treated shCtrl or shMEX3A JHOC5 cells. Cell numbers were counted every day for 3 days. Data are shown as mean ± SD with *P* value based on two-way ANOVA-test (n = 3). ****, P < 0.0001. The experiments were repeated 3 times.

**Table S1.** Short tandem repeat of ovarian cancer cell lines

| OC subtype | OCCC | | | | | HGSOC | |
|---|---|---|---|---|---|---|---|
| Cell Line \STR | TOV21G | JHOC9 | JHOC5 | RMG1 | OVCA429 | KURAMOCHI | OVSAHO |
| TH01 | 8,11 | 6,9 | 7,7 | 6,7 | 9,9 | 6,7 | 6,6 |
| D5S818 | 12,13 | 10,10 | 10,10 | 12,12 | 12,12 | 12,12 | 12,13 |
| D13S317 | 11,12 | 10,10 | 9,12 | 8,12 | 9,12 | 8,12 | 8,8 |
| D7S820 | 12,12 | 8,12 | 12,12 | 11,11 | 10,11 | 11,11 | 8,10 |
| D16S539 | 10,12 | 12,12 | 11,13 | 9,10 | 10,10 | 9,10 | 9,9 |
| CSF1PO | 13,15 | 10,13 | 10,12 | 10,10 | 11,12 | 10,10 | 10,12 |
| Amelogenin | X,X | X,X | X,X | X,X | X,X | X,X | X,X |
| VWA | 17,17 | 17,17 | 14,16 | 17,18 | 16,19 | 17,18 | 14,14 |
| TPOX | 8,11 | 8,11 | 11,11 | 11,11 | 8,12 | 11,11 | 8,11 |
| Match to Test Sample | 100% | 94% | 100% | 94% | 100% | 100% | 100% |
| Database | DSMZ | DSMZ | ExPASy | DSMZ | ExPASy | DSMZ | ExPASy |

**Table S2.** Antibodies.

| Antibodies for immunoblotting (IB) | | | | |
|---|---|---|---|---|
| **Antibody** | **Supplier** | **Cat./RRID** | **Isotype** | **Dilution** |
| GAPDH (D16H11) XP | Cell Signaling | 5174<br>RRID: AB_10622025 | Rabbit IgG | 1:20000 |
| COX IV [Mitochondrial Marker Antibody Sampler Kit] | Cell Signaling | 4850T<br>RRID: AB_2085424 | Rabbit IgG | 1:1000 |
| p53 | Santa Cruz | sc-126<br>RRID: AB_628082 | Mouse monoclonal IgG2a, kappa | 1:5000 |
| Total OXPHOS Human WB Antibody Cocktail | Abcam | ab110411<br>RRID: AB_2756818 | Mouse monoclonal antibodies | 1:1000 |
| TOM20 (F-10) | Santa Cruz | sc-17764<br>RRID: AB_628381 | Mouse monoclonal IgG2a, kappa | 1:5000 |
| SQSTM1/p62 | Abclonal | A19700<br>RRID: AB_2862742 | Rabbit IgG | 1:2000 |
| MEX3A | GeneTex | PJ90803 | Rabbit | 1:10000 |
| DRP1 (D6C7) | Cell Signaling | 8570S<br>RRID: AB_10950498 | Rabbit IgG | 1:5000 |
| MFF | Proteintech | 17090-1-AP<br>RRID: AB_2142463 | Rabbit IgG | 1:5000 |
| PINK1 | Novus | BC100-494<br>RRID: AB_10127658 | Rabbit | 1:2000 |
| PARKIN (D4Z1W) | Cell Signaling | 32833S<br>RRID: AB_3073958 | Rabbit IgG | 1:2000 |
| Phospho-PINK1 (Ser228) (E9K3K) | Cell Signaling | 89010S | Rabbit IgG | 1:2000 |
| Phospho-DRP1 (Ser616) | Cell Signaling | #3455<br>RRID: AB_ 2085352 | Rabbit IgG | 1:5000 |
| LC3B (D11) | Cell Signaling | 3868S<br>RRID: AB_2137707 | Rabbit IgG | 1:1000 |
| **Antibodies for immunofluorescence (IF)** | | | | |
| **Antibody** | **Supplier** | **Cat./RRID** | **Isotype** | **Dilution** |
| TOM20 (F-10) | Santa Cruz | sc-17764<br>RRID: AB_628381 | Mouse monoclonal IgG2a, kappa | 1:50 |
| MFF | Proteintech | 17090-1-AP<br>RRID: AB_2142463 | Rabbit IgG | 1:50 |
| Phospho-DRP1 (Ser616) | Cell Signaling | #3455<br>RRID: AB_2085352 | Rabbit IgG | 1:50 |

**Table S3.** Primers for qPCR

| Gene | Forward (5′→3′) | Reverse (5′→3′) |
|---|---|---|
| MEX3A | GGCAGGCAAGGCTGCAAGA | GGGAGGCACGGATCATGGAG |
| NDUFS1 | GAGTGGACTCTGACACCTTATGC | ACAACATCTGCCTCTTCCACACC |
| NDUFS2 | TACCAAGTTCCTCCAGGAGCCA | GGCAAAACCAGGAGCCTTGATC |
| NDUFB7 | CTGCTCAAGTGCAAGCGTGACA | CGCTCAAACTCCTTCATGCGCA |
| NDUFC2 | AAGCCTTCTGTGGTGCTGTA | ACAGGGTGAAAGGCTGGTTA |
| PGC1A | CCAAAGGATGCGCTCTCGTTCA | CGGTGTCTGTAGTGGCTTGACT |
| NRF1 | GGCAACAGTAGCCACATTGGCT | GTCGTCTGGATGGTCATCTCAC |
| NRF2 | CTGCTGCACTGGAAGGCTATAG | GGTGAGGTCTATATCGGTCATGC |
| TFAM | GTGGTTTTCATCTGTCTTGGCAAG | TTCCCTCCAACGCTGGGCAATT |
| Cyclophilin | GGGTTCCTCCTTTCACAGAATTATT | TTGCCACCAGTGCCATTATG |

**Table S4.** RNAseq: Gene Set Enrichment Analysis (GSEA) on the ranked gene list, determined by the Log Fold-Change. (xlsx file)

**Table S5.** RNAseq: Expression fold-change of the enriched genes in the GSEA-OXPHOS hallmark between shCtrl and shMEX3A TOV21G cells. (xlsx file)

| | ID | Descriptic | setSize | enrichme | NES | pvalue | p.adjust | qvalue | rank | leading_e | core_enri | symbol |
|---|---|---|---|---|---|---|---|---|---|---|---|---|
| HALLMARK_E2F_TARGETS | HALLMAR | HALLMAR | 198 | 0.745362 | 2.021266 | 1.00E-10 | 5.00E-09 | 3.37E-09 | 2197 | tags=54% | 5931/4001 | RBBP7/LMNB1/CDK1/PAICS/TOP2A/POLD3/RACGAP1/ATAD2/HMGB3/BUB1B/PTTG1/AURKA/CDC20/RRM2/KIF4A/USP1/DLGAP5/TACC3/SPAG5/CDCA8/MKI67/MELK/SMC4/ESPL1/BRCA1/BIRC5/MXD3/DEPDC1/PLK1/CCNB2/MCM6/MSH2/E2F8/TMPO/KIF2C/CCP110/MCM4/HMMR/RNASEH2A/KIF22/PMS2/MCM3/PSMC3IP/CSE1L/CENPE/HMGB2/RAD21/ANP32E/CDKN2C/CDCA3/CHEK1/EZH2/DUT/CDKN3/MTHFD2/UBE2S/NBN/SPC25/MCM5/MCM7/TK1/TRA2B/SSRP1/BARD1/SLBP/BRCA2/SMC3/NUP205/NASP/CIT/RAD50/CKS2/CDKN1B/STMN1/RPA3/ASF1A/UBR7/SYNCRIP/PRDX4/DCLRE1B/RAD1/H2AZ1/NOP56/RAD51AP1/ASF1B/RAN/NAA38/RFC1/STAG1/DSCC1/PLK4/CTCF/UBE2T/PPP1R8/RANBP1/HELLS/MYC/SMC1A/TCF19/PHF5A/NUP153/IPO7/NCAPD2/RFC3/PRIM2/SMC6 |
| HALLMARK_TNFA_SIGNALING_VIA_NFKB | HALLMAR | HALLMAR | 179 | -0.7103 | -1.8161 | 5.97E-07 | 1.49E-05 | 1.01E-05 | 1530 | tags=32% | 10611/329 | PDLIM5/BIRC2/F3/SPHK1/BTG3/IRF1/CLCF1/RNF19B/CXCL3/GADD45A/PER1/IER2/EHD1/FJX1/CCNL1/NFKB2/BCL3/FOSL2/G0S2/SERPINB8/NINJ1/KYNU/ZC3H12A/PLAUR/ICAM1/NR4A2/DUSP4/NFKB1/CCND1/LAMB3/DUSP5/PFKFB3/BTG2/DRAM1/CCN1/JUN/TNFAIP2/TNC/CDKN1A/LITAF/INHBA/SNN/CXCL1/RHOB/MAP2K3/CSF1/PLAU/LIF/SAT1/SLC16A6/FOSL1/SLC2A3/DUSP1/IL6/KDM6B/PMEPA1/SERPINE1 |
| HALLMARK_G2M_CHECKPOINT | HALLMAR | HALLMAR | 196 | 0.644246 | 1.74363 | 1.37E-05 | 0.000228 | 0.000154 | 1932 | tags=46% | 1063/4001 | CENPF/LMNB1/E2F2/CDK1/TOP2A/NUP50/KIF11/RACGAP1/NUSAP1/HMGB3/CDC6/EXO1/KIF23/YTHDC1/PTTG1/AURKA/CDC20/KIF4A/STIL/TPX2/KNL1/NEK2/TACC3/HOXC10/KIF5B/MKI67/SMC4/ESPL1/BIRC5/NOTCH2/CCNA2/NCL/PLK1/CCNB2/TTK/MCM6/TMPO/KIF2C/FBXO5/RPS6KA5/HMMR/KIF22/PRC1/H2AZ2/MCM3/SLC7A1/NDC80/CENPE/KIF20B/RAD21/DTYMK/UBE2C/CDKN2C/CHEK1/EZH2/CDC45/PBK/CDKN3/UBE2S/MCM5/SQLE/RAD54L/KIF15/CENPA/TRA2B/RASAL2/FANCC/BUB1/BARD1/BRCA2/DBF4/NASP/CKS2/CDKN1B/STMN1/MAPK14/SYNCRIP/BUB3/H2AZ1/INCENP/CDC7/SMC2/KPNB1/CDC27/CCNF/STAG1/POLQ/EFNA5/PLK4/CTCF |
| HALLMARK_MYC_TARGETS_V1 | HALLMAR | HALLMAR | 199 | 0.616941 | 1.676336 | 3.20E-05 | 0.0004 | 0.000269 | 2767 | tags=38% | 8894/5107 | EIF2S2/HDDC2/RPL22/PTGES3/STARD7/CDC20/VBP1/USP1/NCBP2/HNRNPA1/CCNA2/SRSF3/PHB2/MCM6/PRPS2/NPM1/MCM4/HNRNPR/RUVBL2/RACK1/RPL14/CCT5/RSL1D1/RRM1/MRPS18B/HNRNPA2B1/CDC45/DUT/IMPDH2/SERBP1/MCM5/MCM7/PSMC4/TRA2B/PPIA/CANX/TYMS/MRPL9/NHP2/SYNCRIP/SRSF7/BUB3/PRDX4/H2AZ1/NOP56/KPNB1/RAN/EIF4H/ILF2/FBL/DDX18/RANBP1/MYC/YWHAE/GOT2/CCT3/SNRPA/PWP1/DHX15/IFRD1/POLE3/MAD2L1/ETF1/SSBP1/XPO1/PPM1G/POLD2/FAM120A/SRSF1/GLO1/CSTF2/HNRNPC/DEK/PSMA4/VDAC1 |
| HALLMARK_INTERFERON_ALPHA_RESPONSE | HALLMAR | HALLMAR | 90 | -0.71369 | -1.67632 | 0.000388 | 0.003237 | 0.002181 | 2143 | tags=34% | 64135/569 | IFIH1/PSMB9/HLA-C/IFI27/ELF1/HERC6/PLSCR1/TRAFD1/EIF2AK2/STAT2/PARP12/TRIM5/IFITM2/B2M/IRF9/IRF1/WARS1/RNF31/TRIM14/MOV10/ISG15/IFITM3/UBE2L6/UBA7/TXNIP/OAS1/CD74/CSF1/HELZ2/LGALS3BP/MX1 |
| HALLMARK_P53_PATHWAY | HALLMAR | HALLMAR | 183 | -0.6249 | -1.59724 | 0.00036 | 0.003237 | 0.002181 | 1835 | tags=32% | 351/23479 | APP/ISCU/RB1/IER3/TM7SF3/NOTCH1/ANKRA2/ZMAT3/ADA/SPHK1/RALGDS/IRAK1/RNF19B/CCND3/GADD45A/BAK1/UPP1/APAF1/HSPA4L/ABCC5/S100A10/NINJ1/CD82/SESN1/MAPKAPK3/CCNK/SLC35D1/ITGB4/ZFP36L1/DGKA/EI24/SEC61A1/H1-2/EPS8L2/FDXR/TXNIP/AEN/PHLDA3/S100A4/TM4SF1/POLH/BTG2/BLCAP/DRAM1/TRAF4/TP53/CYFIP2/JUN/BAX/PIDD1/CDKN1A/CCND2/POM121/PLXNB2/MDM2/LIF/SAT1/MKNK2/TGFA |
| HALLMARK_HYPOXIA | HALLMAR | HALLMAR | 178 | -0.62199 | -1.58881 | 0.000614 | 0.004389 | 0.002957 | 1789 | tags=25% | 55577/967 | NAGK/SDC3/IER3/P4HA1/DCN/GPI/KLHL24/ADM/F3/CXCR4/CCN2/CCNG2/SLC6A6/SRPX/MYH9/COL5A1/FOSL2/ERO1A/P4HA2/HSPA5/B3GALT6/PLAUR/GAA/GYS1/VHL/WSB1/ANGPTL4/S100A4/PFKFB3/TGFBI/NDST1/CCN1/JUN/AKAP12/CDKN1A/IDS/MT2A/TGM2/STC1/SLC2A3/DUSP1/IGFBP3/IL6/SERPINE1 |
| HALLMARK_INTERFERON_GAMMA_RESPONSE | HALLMAR | HALLMAR | 164 | -0.62229 | -1.5734 | 0.00091 | 0.005686 | 0.003831 | 2189 | tags=29% | 80830/310 | APOL6/HLA-B/IFIH1/PTPN1/PSMB9/IFI27/HERC6/PLSCR1/TRAFD1/PSMB2/EIF2AK2/STAT2/PARP12/NCOA3/HIF1A/RBCK1/IFITM2/B2M/IRF9/IRF1/WARS1/RNF31/TRIM14/UPP1/OAS2/ISG15/CASP4/RNF213/IFITM3/STAT3/UBE2L6/ICAM1/NFKB1/TXNIP/MVP/ZNFX1/IFNAR2/TNFAIP2/OAS3/CDKN1A/MT2A/CD74/PML/HELZ2/IL6/LGALS3BP/MX1/TAPBP |
| HALLMARK_EPITHELIAL_MESENCHYMAL_TRANSITIC | HALLMAR | HALLMAR | 176 | -0.60068 | -1.53699 | 0.001406 | 0.007809 | 0.005261 | 1094 | tags=26% | 10398/219 | MYL9/FBLN2/COPA/COL5A1/SFRP1/WIPF1/LOXL2/COL11A1/SERPINH1/LRP1/PLAUR/TAGLN/CAP2/NNMT/FLNA/ITGB1/TNFRSF12A/FBN1/IGFBP4/ANPEP/THBS2/TGFBI/COL4A1/CCN1/JUN/TNC/INHBA/COL4A2/NID2/ITGA5/CXCL1/RHOB/COL1A2/IL32/BDNF/TGM2/SAT1/COL7A1/ACTA2/COL1A1/IGFBP3/IL6/NT5E/PMEPA1/SERPINE1/TFPI2 |
| HALLMARK_COMPLEMENT | HALLMAR | HALLMAR | 156 | -0.58627 | -1.47136 | 0.003465 | 0.017324 | 0.011671 | 1471 | tags=24% | 27445/306 | PCLO/KCNIP2/CD59/CASP10/PCSK9/ITGAM/RCE1/CASP3/SRC/CP/PSMB9/DYRK2/PLSCR1/ERAP2/SH2B3/PIK3R5/F3/IRF1/PRDM4/CBLB/EHD1/CALM3/KYNU/CLU/CASP4/HSPA5/LRP1/PLAUR/DUSP5/BRPF3/COL4A2/C3/CXCL1/GNAI2/IL6/SERPINE1/TFPI2 |
| HALLMARK_ESTROGEN_RESPONSE_EARLY | HALLMAR | HALLMAR | 174 | -0.57418 | -1.46556 | 0.005479 | 0.022831 | 0.015381 | 1342 | tags=22% | 5029/1154 | P2RY2/CISH/OPN3/PPIF/CANT1/FRK/HSPB8/SVIL/MYBBP1A/ITPK1/NAV2/SCNN1A/ADCY9/FLNB/ALDH3B1/GREB1/RAPGEFL1/UGCG/KRT8/SLC37A1/CELSR2/WWC1/CCND1/PODXL/KRT18/IGFBP4/PTGES/THSD4/ELF3/ABHD2/TSKU/RAB17/KAZN/SLC7A5/MED24/FASN/TGM2/SLC26A2/NCOR2 |
| HALLMARK_IL2_STAT5_SIGNALING | HALLMAR | HALLMAR | 158 | -0.58289 | -1.46306 | 0.005303 | 0.022831 | 0.015381 | 1408 | tags=16% | 4641/896/ | MYO1C/CCND3/CISH/BMPR2/MAPKAPK2/IGF2R/ALCAM/ODC1/NRP1/IFITM3/SOCS2/ADAM19/PLEC/ETV4/F2RL2/SPRY4/NCS1/RHOB/FURIN/EMP1/CSF1/CCND2/LIF/TGM2/SLC2A3/NT5E |
| HALLMARK_APICAL_JUNCTION | HALLMAR | HALLMAR | 161 | -0.58094 | -1.46178 | 0.006319 | 0.024304 | 0.016373 | 1763 | tags=29% | 9672/8189 | SDC3/SYMPK/INPPL1/ICAM2/NEGR1/PLCG1/MAP4K2/SPEG/CD276/NLGN2/DLG1/TNFRSF11B/ACTN4/TMEM8B/CERCAM/NECTIN2/PTK2/CLDN15/MYL9/AKT2/MYH9/IRS1/SORBS3/NECTIN1/NF1/TAOK2/ITGB4/NECTIN4/VASP/ICAM1/MAPK13/COL17A1/LAMB3/RRAS/ITGB1/CNN2/FBN1/TGFBI/JUP/PKD1/ZYX/FLNC/GNAI2/ITGA3/ACTN1/FSCN1/NF2 |
| HALLMARK_OXIDATIVE_PHOSPHORYLATION | HALLMAR | HALLMAR | 198 | 0.503654 | 1.365804 | 0.011342 | 0.040506 | 0.027288 | 2711 | tags=31%, | 481/509/3 | ATP1B1/ATP5F1C/ACAA1/COX7A2L/CYCS/PDK4/SDHD/LDHB/PDHB/ALDH6A1/BDH2/PHB2/IDH2/COX15/MFN2/HTRA2/ECH1/IDH3B/MDH1/ECHS1/SLC25A5/TIMM13/MRPL15/ATP6V0E1/MRPL35/MRPS15/MTRR/MPC1/SDHC/ACAT1/ATP5PF/ATP6V0B/DLD/NQO2/NDUFS2/PDHX/NDUFS1/GOT2/ATP5PD/GRPEL1/TIMM9/GPX4/UQCRFS1/DLST/POR/ATP5F1A/UQCRB/NDUFB7/ATP5ME/TOMM22/NDUFA9/RETSAT/TIMM10/CYB5A/MRPS12/GLUD1/ECI1/TIMM17A/ACAA2/CASP7/NDUFC2 |
| HALLMARK_APOPTOSIS | HALLMAR | HALLMAR | 143 | -0.57781 | -1.43455 | 0.015366 | 0.051219 | 0.034506 | 1430 | tags=23% | 10950/365 | BTG3/IRF1/LMNA/SMAD7/CTNNB1/GADD45A/IGFBP6/PTK2/IGF2R/SPTAN1/GSN/CLU/CASP4/IFITM3/DAP/PLCB2/ERBB2/EREG/FDXR/CCND1/TXNIP/SLC20A1/TNFRSF12A/KRT18/BTG2/JUN/BAX/CDKN1A/RHOB/EMP1/CCND2/SAT1/IL6 |
| HALLMARK_ALLOGRAFT_REJECTION | HALLMAR | HALLMAR | 126 | -0.57595 | -1.40381 | 0.026329 | 0.082279 | 0.05543 | 1355 | tags=15% | 7453/896/ | WARS1/CCND3/TLR6/ITGB2/BCL3/GALNT1/AKT1/ICAM1/EREG/FLNA/SRGN/IFNAR2/INHBA/CD74/CSF1/CCND2/LIF/IL6/TAPBP |
| HALLMARK_KRAS_SIGNALING_UP | HALLMAR | HALLMAR | 139 | -0.54504 | -1.34697 | 0.030361 | 0.089297 | 0.060158 | 1083 | tags=19% | 208/26499 | AKT2/PLEK2/SPRY2/ITGB2/G0S2/ERO1A/TFPI/SOX9/NRP1/MMP11/PLAUR/EREG/EPHB2/ANGPTL4/AKAP12/ETV4/JUP/GNG11/INHBA/EMP1/EPB41L3/DCBLD2/CCND2/PLAU/LIF/IGFBP3 |
| HALLMARK_INFLAMMATORY_RESPONSE | HALLMAR | HALLMAR | 142 | -0.5437 | -1.34818 | 0.040108 | 0.111412 | 0.075056 | 1738 | tags=20% | 3091/7855 | HIF1A/FZD5/PIK3R5/ADM/ADRM1/F3/SPHK1/EBI3/IRF1/P2RY2/MET/ADGRE1/CD82/AXL/PLAUR/ICAM1/EREG/NFKB1/BTG2/CDKN1A/INHBA/ITGA5/DCBLD2/CSF1/LIF/IL6/SERPINE1/TAPBP |
| HALLMARK_TGF_BETA_SIGNALING | HALLMAR | HALLMAR | 53 | -0.62632 | -1.36498 | 0.051786 | 0.136278 | 0.091808 | 1298 | tags=28% | 3065/4097 | HDAC1/SMAD7/CTNNB1/ENG/BMPR2/SMURF2/SMURF1/LTBP2/BCAR3/SLC20A1/SKI/FURIN/NCOR2/PMEPA1/SERPINE1 |
| HALLMARK_KRAS_SIGNALING_DN | HALLMAR | HALLMAR | 107 | -0.53892 | -1.29256 | 0.074627 | 0.177683 | 0.119702 | 1014 | tags=11% | 54842/974 | MFSD6/CLSTN3/GPR3/NR4A2/CELSR2/SPTBN2/BTG2/KMT2D/SPHK2/SYNPO/SNN/MX1 |
| HALLMARK_MTORC1_SIGNALING | HALLMAR | HALLMAR | 194 | 0.447796 | 1.214949 | 0.073314 | 0.177683 | 0.119702 | 1242 | tags=21% | 112399/88 | EGLN3/EIF2S2/CD9/SCD/HMGCS1/FADS1/AURKA/DHFR/RRM2/NUPR1/PSAT1/PSME3/CYB5B/USO1/IGFBP5/TMEM97/PLK1/SKAP2/MCM4/SQSTM1/DDIT3/VLDLR/GGA2/CCNG1/MTHFD2/SQLE/NFIL3/SLC2A1/SLC1A4/CTSC/PITPNB/PSMC4/TPI1/EBP/PPIA/BUB1/CANX/SLC37A4/NUP205/IFI30 |
| HALLMARK_HEDGEHOG_SIGNALING | HALLMAR | HALLMAR | 33 | -0.64566 | -1.2847 | 0.125455 | 0.258427 | 0.174098 | 1009 | tags=27% | 4627/9289 | MYH9/ADGRG1/TLE1/NF1/NRP1/SHH/NRP2/L1CAM/PML |
| HALLMARK_PEROXISOME | HALLMAR | HALLMAR | 92 | 0.497411 | 1.218032 | 0.118227 | 0.258427 | 0.174098 | 963 | tags=12% | 30/216/71 | ACAA1/ALDH1A1/TOP2A/FADS1/EPHX2/MSH2/HSD17B11/FDPS/ATXN1/IDH2/ECH1 |
| HALLMARK_UV_RESPONSE_DN | HALLMAR | HALLMAR | 138 | -0.48578 | -1.20166 | 0.122186 | 0.258427 | 0.174098 | 1530 | tags=22% | 10611/173 | PDLIM5/DLG1/F3/PTPRM/PPARG/SMAD7/MET/IRS1/PIK3CD/TFPI/COL11A1/TENT4A/APBB2/RUNX1/NRP1/ERBB2/CAP2/NFKB1/PIAS3/RBPMS/MTA1/CCN1/ABCC1/FZD2/PTPN21/CELF2/COL1A2/BDNF/DUSP1/COL1A1/SERPINE1 |
| HALLMARK_ESTROGEN_RESPONSE_LATE | HALLMAR | HALLMAR | 168 | 0.438529 | 1.169039 | 0.129213 | 0.258427 | 0.174098 | 1309 | tags=20% | 1152/1363 | CKB/CPE/IGSF1/CD9/KIF20A/TOP2A/IMPA2/ID2/CDC6/LAMC2/CDC20/STIL/TPD52L1/CXCL12/SGK1/SFN/EMP2/IL17RB/BCL2/METTL3/RNASEH2A/IDH2/HOMER2/ARL3/ALDH3A2/PRSS23/RBBP8/MOCS2/SLC1A4/MAPT/PRKAR2B/FABP5/CHPT1 |
| HALLMARK_IL6_JAK_STAT3_SIGNALING | HALLMAR | HALLMAR | 59 | -0.5548 | -1.24055 | 0.137631 | 0.264674 | 0.178307 | 2134 | tags=29% | 7040/5770 | TGFB1/PTPN1/CBL/STAT2/PIK3R5/TYK2/IRF9/EBI3/IRF1/CXCL3/BAK1/STAT3/TNFRSF12A/JUN/CXCL1/CSF1/IL6 |
| HALLMARK_MITOTIC_SPINDLE | HALLMAR | HALLMAR | 199 | 0.418765 | 1.137859 | 0.148368 | 0.274755 | 0.185098 | 1664 | tags=29% | 1063/4001 | CENPF/LMNB1/CDK1/TOP2A/KIF11/RACGAP1/NUSAP1/KIF23/AURKA/KIF4A/DLGAP5/TPX2/NEK2/WASF1/CNTRL/KIF5B/SMC4/ESPL1/TBCD/TLK1/BIRC5/NOTCH2/PLK1/CCNB2/TTK/KIF2C/FBXO5/KIF22/PRC1/KIF3B/MYO1E/CCDC88A/NDC80/CENPE/KIF20B/CDC42EP4/ARHGEF3/PCM1/ROCK1/ARF6/KIF15/RASAL2/BUB1/BRCA2/SMC3/ARHGAP29/ANLN/RAPGEF5/EPB41/NIN/MYH10/CEP57/INCENP/APC/PCNT/KNTC1/PPP4R2/PKD2 |
| HALLMARK_COAGULATION | HALLMAR | HALLMAR | 94 | -0.49225 | -1.15831 | 0.198642 | 0.354717 | 0.238968 | 1243 | tags=18% | 726/2152/ | CAPN5/F3/CFI/SPARC/CAPN2/GSN/CLU/LRP1/MMP11/FBN1/RAPGEF3/F2RL2/C3/FURIN/PLAU/SERPINE1/TFPI2 |
| HALLMARK_MYOGENESIS | HALLMAR | HALLMAR | 141 | -0.45437 | -1.12579 | 0.216693 | 0.373609 | 0.251695 | 1187 | tags=21% | 1760/6675 | DMPK/SPARC/AKT2/SMTN/PC/HSPB8/SVIL/TSC2/NAV2/MYH9/SORBS3/ADCY9/SPTAN1/GSN/CLU/ITGB4/MAPRE3/GAA/TAGLN/ITGB1/MEF2D/KCNH1/KIFC3/AGRN/CDKN1A/COL4A2/PLXNB2/COL1A1/IGFBP3 |
| HALLMARK_BILE_ACID_METABOLISM | HALLMAR | HALLMAR | 79 | 0.455174 | 1.09517 | 0.27685 | 0.432578 | 0.291421 | 1846 | tags=25% | 1593/216/ | CYP27A1/ALDH1A1/FADS1/EPHX2/HACL1/HSD17B11/ATXN1/IDH2/EFHC1/TFCP2L1/PXMP2/SLC35B2/PEX12/SERPINA6/SLC23A1/CAT/SLCO1A2/GCLM/PFKM/NPC1 |
| HALLMARK_XENOBIOTIC_METABOLISM | HALLMAR | HALLMAR | 151 | 0.41651 | 1.090396 | 0.259777 | 0.432578 | 0.291421 | 1141 | tags=15% | 1593/6747 | CYP27A1/SSR3/NQO1/PTGES3/AKR1C3/PDK4/ID2/LPIN2/TMBIM6/ENTPD5/CNDP2/TMEM97/HACL1/FBLN1/HMOX1/CBR1/DDAH2/ETS2/GCNT2/ECH1/RAP1GAP/PTGDS/EPHX1 |
| HALLMARK_FATTY_ACID_METABOLISM | HALLMAR | HALLMAR | 137 | 0.412561 | 1.079277 | 0.275591 | 0.432578 | 0.291421 | 1236 | tags=17% | 30/8801/2 | ACAA1/SUCLG2/ALDH1A1/HMGCS1/SDHD/PDHB/OSTC/CRYZ/GLUL/CBR1/HSD17B11/LGALS1/HADH/ALDH3A2/ECH1/ACAT2/NBN/ENO2/IDH3B/MDH1/MGLL/EPHX1/ECHS1 |
| HALLMARK_MYC_TARGETS_V2 | HALLMAR | HALLMAR | 58 | 0.47082 | 1.081311 | 0.316159 | 0.479029 | 0.322714 | 2358 | tags=24% | 27346/534 | TMEM97/PLK1/NPM1/MCM4/MCM5/NOP56/RCL1/PLK4/DDX18/NDUFAF4/MYC/NOP2/UNG/MRTO4 |
| HALLMARK_CHOLESTEROL_HOMEOSTASIS | HALLMAR | HALLMAR | 71 | 0.448571 | 1.065737 | 0.339667 | 0.485239 | 0.326898 | 1305 | tags=25% | 928/6319/ | CD9/SCD/HMGCS1/GLDC/ANXA13/ACTG1/ANTXR2/TMEM97/FDPS/JAG1/ECH1/ACAT2/SQLE/NFIL3/PMVK/EBP/ERRFI1/FABP5 |
| HALLMARK_SPERMATOGENESIS | HALLMAR | HALLMAR | 73 | 0.444859 | 1.06318 | 0.332536 | 0.485239 | 0.326898 | 1803 | tags=30% | 983/6790/ | CDK1/AURKA/NEK2/TNNI3/CSNK2A2/CCNB2/TTK/KIF2C/YBX2/GSTM3/NCAPH/EZH2/CDKN3/PARP2/CAMK4/BUB1/DBF4/NPHP1/TSN/SIRT1/PCSK1N/MAP7 |
| HALLMARK_ANDROGEN_RESPONSE | HALLMAR | HALLMAR | 93 | -0.44584 | -1.04874 | 0.365482 | 0.507614 | 0.341972 | 646 | tags=13% | 6722/207/ | SRF/AKT1/KRT8/CCND1/ABHD2/AKAP12/DHCR24/PTPN21/SAT1/SLC26A2/ACTN1/PMEPA1 |
| HALLMARK_NOTCH_SIGNALING | HALLMAR | HALLMAR | 30 | 0.496554 | 1.030875 | 0.413567 | 0.558874 | 0.376505 | 1568 | tags=27% | 4854/5578 | NOTCH3/PRKCA/NOTCH2/JAG1/SKP1/LFNG/PSENEN/DTX4 |
| HALLMARK_WNT_BETA_CATENIN_SIGNALING | HALLMAR | HALLMAR | 38 | -0.50097 | -1.02451 | 0.435943 | 0.558901 | 0.376523 | 1718 | tags=24% | 4851/1001 | NOTCH1/HDAC5/PPARD/CTNNB1/SKP2/MAML1/TP53/CCND2/NCOR2 |
| HALLMARK_GLYCOLYSIS | HALLMAR | HALLMAR | 185 | -0.39402 | -1.00656 | 0.431853 | 0.558901 | 0.376523 | 1230 | tags=12% | 54480/423 | CHPF2/MET/PYGB/PC/COL5A1/SLC25A13/ERO1A/SOX9/P4HA2/HSPA5/B3GALT6/AK3/GYS1/ANGPTL4/TGFBI/ELF3/AGRN/MED24/STC1/IGFBP3/NT5E/SLC16A3/TGFA |
| HALLMARK_PROTEIN_SECRETION | HALLMAR | HALLMAR | 93 | 0.404494 | 0.996211 | 0.454988 | 0.568735 | 0.383148 | 1899 | tags=22% | 11079/107 | RER1/ADAM10/USO1/SCAMP1/AP2M1/ATP1A1/EGFR/SEC22B/AP3B1/ANP32E/GOLGA4/PPT1/CTSC/TMED10/SCRN1/GOSR2/RAB2A/CLTA/RPS6KA3/SNX2 |
| HALLMARK_DNA_REPAIR | HALLMAR | HALLMAR | 147 | 0.376199 | 0.98461 | 0.480874 | 0.584244 | 0.393596 | 1689 | tags=20% | 51292/258 | GMPR2/MPC2/POLD3/SMAD5/NCBP2/FEN1/TAF13/CETN2/ERCC4/POLR2G/ZWINT/POLR2D/REV3L/RFC5/DUT/IMPDH2/PRIM1/NELFE/NT5C/SSRP1/POLR1D/MRPL40/TYMS/CSTF3/NME4/RPA3/GTF2F1/UPF3B/PDE4B/RAD51 |
| HALLMARK_UV_RESPONSE_UP | HALLMAR | HALLMAR | 141 | 0.373544 | 0.977663 | 0.490765 | 0.584244 | 0.393596 | 1340 | tags=14% | 682/30/54 | BSG/ACAA1/PPAT/GPX3/TYRO3/MAPK8IP2/TMBIM6/FEN1/CYB5B/GGH/SQSTM1/HMOX1/CTSV/PARP2/ENO2/BTG1/PPT1/EPHX1/CHRNA5/PLCL1 |
| HALLMARK_ANGIOGENESIS | HALLMAR | HALLMAR | 27 | -0.50188 | -0.97214 | 0.525362 | 0.610886 | 0.411545 | 744 | tags=30% | 3685/7410 | ITGAV/VAV2/APP/PTK2/NRP1/S100A4/CCND2/STC1 |
| HALLMARK_PI3K_AKT_MTOR_SIGNALING | HALLMAR | HALLMAR | 95 | -0.37685 | -0.88772 | 0.69322 | 0.761431 | 0.512964 | 1037 | tags=24% | 10097/291 | ACTR2/TBK1/ACACA/RAC1/STAT2/PTEN/MAPKAP1/ARF1/PLCG1/CXCR4/RPTOR/TSC2/AKT1S1/RPS6KA1/AKT1/PPP2R1B/ARHGDIA/SMAD2/CDKN1A/GRK2/PAK4/MAP2K3/MKNK2 |
| HALLMARK_UNFOLDED_PROTEIN_RESPONSE | HALLMAR | HALLMAR | 109 | -0.36422 | -0.87633 | 0.725914 | 0.761431 | 0.512964 | 1954 | tags=18% | 6464/4800 | SHC1/NFYA/SPCS3/DCTN1/HYOU1/EDC4/KHSRP/DCP2/TATDN2/SEC31A/ERO1A/HSPA5/EDEM1/SRPRA/ARFGAP1/ATP6V0D1/EIF2AK3/EIF4G1/ALDH18A1/SLC7A5 |
| HALLMARK_REACTIVE_OXYGEN_SPECIES_PATHWAY | HALLMAR | HALLMAR | 45 | 0.392609 | 0.863814 | 0.70615 | 0.761431 | 0.512964 | 2422 | tags=29% | 1728/2878 | NQO1/GPX3/SELENOS/CDKN2D/PRDX4/TXNRD1/CAT/GCLM/NDUFS2/LAMTOR5/GPX4/GCLC/GLRX |
| HALLMARK_PANCREAS_BETA_CELLS | HALLMAR | HALLMAR | 23 | 0.443923 | 0.862539 | 0.679739 | 0.761431 | 0.512964 | 1998 | tags=30% | 5122/1056 | PCSK1/SCGN/SRPRB/SPCS1/IAPP/SEC11A/PAX6 |
| HALLMARK_APICAL_SURFACE | HALLMAR | HALLMAR | 36 | -0.40543 | -0.82367 | 0.730973 | 0.761431 | 0.512964 | 1975 | tags=28% | 50617/351 | ATP6V0A4/APP/NCOA6/CROCC/TMEM8B/ATP8B1/SRPX/PLAUR/EPHB4/DCBLD2 |
| HALLMARK_ADIPOGENESIS | HALLMAR | HALLMAR | 181 | 0.321926 | 0.868452 | 0.815562 | 0.819165 | 0.551859 | 1760 | tags=13% | 10580/753 | SORBS1/YWHAG/ATP1B3/GPX3/FZD4/RMDN3/EPHX2/DBT/MYLK/STOM/HADH/ITGA7/CDKN2C/ECH1/MGLL/CD151/ECHS1/TKT/MRPL15/SLC1A5/ELOVL6/CAT/PPM1B/SDHC |
| HALLMARK_HEME_METABOLISM | HALLMAR | HALLMAR | 167 | -0.33721 | -0.85378 | 0.819165 | 0.819165 | 0.551859 | 1513 | tags=12% | 9091/2336 | PIGQ/ARHGEF12/CCND3/TMCC2/ADIPOR1/CPOX/LRP10/MARCHF2/PC/NARF/MARCHF8/P4HA2/CLCN3/BTG2/RNF123/ATP6V0A1/C3/MAP2K3/SEC14L1/DMTN |

| | shMEX3A_1 TOVshM_1_S97 | shMEX3A_2 TOVshM_2_S98 | shMEX3A_3 TOVshM_3_S99 | shCtrl_1 TOVT2_1_S94 | shCtrl_2 TOVT2_2_S95 | shCtrl_3 TOVT2_3_S96 | shCtrl vs shMEX3A logFC | logCPM | PValue | FDR | DGEtest |
|---|---|---|---|---|---|---|---|---|---|---|---|
| ATP1B1 | 7.0757 | 6.9859 | 6.9924 | 8.6649 | 8.6730 | 8.6998 | 1.6610 | 8.0753 | 0 | 0 | 1 |
| ATP5F1C | 6.2968 | 6.2428 | 6.2253 | 7.9530 | 7.8418 | 7.9197 | 1.6509 | 7.3043 | 0 | 0 | 1 |
| ACAA1 | 4.4018 | 4.3188 | 4.3558 | 5.9334 | 5.9950 | 5.9691 | 1.6093 | 5.3727 | 1.83E-276 | 3.22E-274 | 1 |
| COX7A2L | 5.6974 | 5.8001 | 5.7424 | 7.4610 | 7.2950 | 7.2749 | 1.5996 | 6.7574 | 4.39E-226 | 5.09E-224 | 1 |
| SDHD | 4.4869 | 4.3803 | 4.2789 | 5.6109 | 5.5622 | 5.7209 | 1.2479 | 5.1390 | 6.21E-109 | 2.04E-107 | 1 |
| CYCS | 5.1711 | 5.1348 | 5.2188 | 6.3213 | 6.2164 | 6.3810 | 1.1362 | 5.8490 | 6.18E-128 | 2.65E-126 | 1 |
| NDUFC2 | 1.1246 | 0.6170 | 0.9551 | 2.0272 | 1.8832 | 1.7724 | 1.0135 | 1.4814 | 3.50E-10 | 1.08E-09 | 1 |
| PDHB | 5.9133 | 5.8219 | 5.7723 | 6.8344 | 6.7319 | 6.7722 | 0.9432 | 6.3843 | 2.36E-100 | 6.73E-99 | 0 |
| ALDH6A1 | 4.8632 | 4.7704 | 4.8656 | 5.7320 | 5.8064 | 5.7304 | 0.9256 | 5.3652 | 8.58E-89 | 2.06E-87 | 0 |
| BDH2 | 4.1770 | 4.1805 | 4.2591 | 5.1106 | 5.0506 | 5.1101 | 0.8901 | 4.7132 | 6.15E-66 | 9.78E-65 | 0 |
| LDHB | 9.4360 | 9.4231 | 9.3423 | 10.2316 | 10.1671 | 10.2543 | 0.8170 | 9.8667 | 7.36E-106 | 2.25E-104 | 0 |
| PDK4 | 8.9213 | 8.8999 | 8.8978 | 9.7640 | 9.7178 | 9.6767 | 0.8137 | 9.3697 | 2.71E-122 | 1.07E-120 | 0 |
| HTRA2 | 4.3342 | 4.2799 | 4.3558 | 5.1135 | 4.9710 | 5.0828 | 0.7388 | 4.7356 | 2.26E-41 | 2.09E-40 | 0 |
| PHB2 | 7.8702 | 7.8858 | 7.9653 | 8.5041 | 8.5168 | 8.5151 | 0.6049 | 8.2409 | 1.57E-64 | 2.43E-63 | 0 |
| MPC1 | 3.4744 | 3.5857 | 3.6274 | 4.2281 | 4.1163 | 4.1195 | 0.5973 | 3.8886 | 4.56E-19 | 2.17E-18 | 0 |
| COX15 | 6.4349 | 6.4050 | 6.3861 | 6.9263 | 6.9540 | 7.0298 | 0.5617 | 6.7166 | 7.68E-45 | 7.70E-44 | 0 |
| IDH2 | 6.8193 | 6.9043 | 6.8970 | 7.4116 | 7.4357 | 7.3801 | 0.5354 | 7.1660 | 3.24E-45 | 3.28E-44 | 0 |
| TIMM13 | 5.7644 | 5.7538 | 5.7684 | 6.4117 | 6.1815 | 6.2524 | 0.5248 | 6.0477 | 7.08E-25 | 4.12E-24 | 0 |
| IDH3B | 5.8787 | 6.0102 | 6.0188 | 6.5036 | 6.4921 | 6.4826 | 0.5230 | 6.2546 | 9.21E-34 | 6.94E-33 | 0 |
| ATP5ME | 4.5283 | 4.4727 | 4.6019 | 5.2490 | 4.9501 | 4.9165 | 0.5159 | 4.8136 | 5.25E-12 | 1.80E-11 | 0 |
| MDH1 | 6.6498 | 6.5144 | 6.6093 | 7.0784 | 7.0721 | 7.1560 | 0.5115 | 6.8694 | 4.05E-33 | 3.01E-32 | 0 |
| MFN2 | 7.3295 | 7.3242 | 7.2691 | 7.7614 | 7.8387 | 7.8092 | 0.4949 | 7.5766 | 1.46E-41 | 1.36E-40 | 0 |
| MRPL35 | 4.5963 | 4.5518 | 4.6228 | 5.0894 | 5.0208 | 5.1216 | 0.4912 | 4.8537 | 3.88E-22 | 2.06E-21 | 0 |
| TIMM10 | 3.9035 | 3.9644 | 4.1601 | 4.6205 | 4.4722 | 4.3697 | 0.4848 | 4.2695 | 7.12E-11 | 2.29E-10 | 0 |
| ECH1 | 7.5216 | 7.5501 | 7.5865 | 8.0964 | 8.0178 | 7.9817 | 0.4804 | 7.8126 | 6.54E-36 | 5.24E-35 | 0 |
| TIMM9 | 3.7406 | 3.6984 | 3.5604 | 4.1816 | 4.0831 | 4.1823 | 0.4799 | 3.9308 | 3.03E-13 | 1.12E-12 | 0 |
| UQCRB | 6.9397 | 6.9850 | 6.8546 | 7.5491 | 7.2794 | 7.3237 | 0.4617 | 7.1774 | 2.23E-12 | 7.83E-12 | 0 |
| SLC25A5 | 8.1596 | 8.1488 | 8.2771 | 8.6203 | 8.5765 | 8.6728 | 0.4281 | 8.4256 | 5.79E-26 | 3.50E-25 | 0 |
| NQO2 | 4.9358 | 5.0543 | 5.0922 | 5.5254 | 5.4029 | 5.4015 | 0.4185 | 5.2513 | 1.48E-15 | 6.07E-15 | 0 |
| MRPL15 | 5.5671 | 5.5971 | 5.5622 | 5.9864 | 5.9371 | 6.0204 | 0.4064 | 5.7929 | 2.11E-23 | 1.17E-22 | 0 |
| ECHS1 | 7.2595 | 7.2234 | 7.1853 | 7.6431 | 7.5864 | 7.6525 | 0.4045 | 7.4396 | 5.63E-28 | 3.59E-27 | 0 |
| PDHX | 4.9437 | 4.8436 | 4.7860 | 5.2993 | 5.1987 | 5.2818 | 0.4015 | 5.0746 | 7.22E-15 | 2.87E-14 | 0 |
| SDHC | 6.7918 | 6.6387 | 6.6429 | 7.0485 | 7.0670 | 7.1586 | 0.3992 | 6.9061 | 4.29E-18 | 1.96E-17 | 0 |
| ATP6V0E1 | 6.2249 | 6.1923 | 6.2417 | 6.6597 | 6.5666 | 6.6111 | 0.3949 | 6.4295 | 7.96E-23 | 4.33E-22 | 0 |
| MTRR | 5.5854 | 5.5378 | 5.5549 | 5.8945 | 5.9751 | 5.9902 | 0.3941 | 5.7692 | 5.76E-21 | 2.92E-20 | 0 |
| MRPS15 | 5.5883 | 5.5025 | 5.6013 | 5.9691 | 5.9291 | 5.9668 | 0.3933 | 5.7722 | 5.72E-21 | 2.90E-20 | 0 |
| NDUFB7 | 6.0749 | 6.1554 | 6.2590 | 6.6694 | 6.5545 | 6.4344 | 0.3925 | 6.3731 | 4.71E-12 | 1.62E-11 | 0 |
| ATP5PF | 5.4680 | 5.5690 | 5.5810 | 6.0127 | 5.8732 | 5.8760 | 0.3839 | 5.7438 | 1.13E-15 | 4.65E-15 | 0 |
| ATP6V0B | 5.8119 | 5.7048 | 5.8607 | 6.1262 | 6.1905 | 6.1354 | 0.3592 | 5.9818 | 1.28E-15 | 5.27E-15 | 0 |
| ATP5PD | 6.2253 | 6.3222 | 6.3211 | 6.7304 | 6.5959 | 6.5848 | 0.3494 | 6.4750 | 3.76E-14 | 1.45E-13 | 0 |
| CYB5A | 4.6994 | 4.6549 | 4.8059 | 5.0972 | 5.0132 | 5.0668 | 0.3437 | 4.8988 | 7.38E-11 | 2.37E-10 | 0 |
| GRPEL1 | 5.3211 | 5.4348 | 5.3809 | 5.7019 | 5.6804 | 5.7738 | 0.3394 | 5.5595 | 5.71E-14 | 2.18E-13 | 0 |
| ACAT1 | 7.1545 | 7.2937 | 7.2111 | 7.5805 | 7.5312 | 7.5474 | 0.3320 | 7.3967 | 1.10E-16 | 4.72E-16 | 0 |
| MRPS12 | 5.2226 | 5.2119 | 5.2359 | 5.6535 | 5.4512 | 5.5418 | 0.3307 | 5.3972 | 8.20E-11 | 2.63E-10 | 0 |
| GPX4 | 7.9336 | 7.9930 | 8.0783 | 8.4179 | 8.3045 | 8.2611 | 0.3271 | 8.1752 | 3.48E-13 | 1.28E-12 | 0 |
| UQCRFS1 | 6.2789 | 6.2351 | 6.3045 | 6.6740 | 6.5141 | 6.5919 | 0.3237 | 6.4429 | 3.49E-13 | 1.28E-12 | 0 |
| DLD | 6.5711 | 6.5615 | 6.4620 | 6.8568 | 6.8513 | 6.8403 | 0.3159 | 6.6999 | 1.40E-15 | 5.75E-15 | 0 |
| NDUFS1 | 7.5538 | 7.4523 | 7.4746 | 7.7539 | 7.7929 | 7.8577 | 0.3079 | 7.6562 | 8.50E-15 | 3.37E-14 | 0 |
| NDUFS2 | 7.2658 | 7.2688 | 7.2831 | 7.6103 | 7.5441 | 7.5247 | 0.2881 | 7.4235 | 3.07E-15 | 1.24E-14 | 0 |
| CASP7 | 5.1288 | 5.0589 | 5.1377 | 5.3752 | 5.4230 | 5.3827 | 0.2864 | 5.2569 | 2.88E-10 | 8.93E-10 | 0 |
| TIMM17A | 5.2503 | 5.3405 | 5.3107 | 5.6069 | 5.5326 | 5.6141 | 0.2848 | 5.4502 | 2.10E-10 | 6.57E-10 | 0 |
| RETSAT | 6.5667 | 6.4602 | 6.4817 | 6.7173 | 6.8014 | 6.8328 | 0.2808 | 6.6505 | 6.47E-11 | 2.08E-10 | 0 |
| ECI1 | 5.6085 | 5.6141 | 5.6756 | 5.9187 | 5.8423 | 5.9413 | 0.2709 | 5.7731 | 1.77E-10 | 5.55E-10 | 0 |
| ATP5F1A | 9.2012 | 9.1286 | 9.1352 | 9.3966 | 9.3820 | 9.4948 | 0.2700 | 9.2967 | 1.63E-12 | 5.78E-12 | 0 |
| GOT2 | 7.3490 | 7.3194 | 7.3270 | 7.5779 | 7.5999 | 7.6214 | 0.2680 | 7.4719 | 1.13E-14 | 4.46E-14 | 0 |
| DLST | 7.6755 | 7.7070 | 7.6694 | 7.8890 | 7.9619 | 7.9646 | 0.2544 | 7.8170 | 1.09E-12 | 3.90E-12 | 0 |
| ACAA2 | 6.7021 | 6.6307 | 6.6391 | 6.9151 | 6.8573 | 6.9564 | 0.2530 | 6.7895 | 2.61E-10 | 8.10E-10 | 0 |
| NDUFA9 | 6.5381 | 6.5439 | 6.4807 | 6.8025 | 6.7532 | 6.7626 | 0.2513 | 6.6528 | 5.13E-11 | 1.66E-10 | 0 |
| TOMM22 | 6.9628 | 6.9513 | 6.9820 | 7.2460 | 7.1689 | 7.2102 | 0.2442 | 7.0922 | 3.75E-11 | 1.23E-10 | 0 |
| POR | 8.0006 | 8.0623 | 8.0422 | 8.2878 | 8.2873 | 8.2585 | 0.2426 | 8.1616 | 1.15E-12 | 4.11E-12 | 0 |
| GLUD1 | 8.0072 | 7.9763 | 7.9557 | 8.1660 | 8.2065 | 8.2500 | 0.2278 | 8.0984 | 1.58E-10 | 4.99E-10 | 0 |
